# Central amygdalar PKCδ neurons mediate fentanyl withdrawal

**DOI:** 10.1101/2025.02.21.639538

**Authors:** Lisa M. Wooldridge, Jacqueline W.K. Wu, Adrienne Y. Jo, Morgan Zinn, Angela M. Lee, Malaika Mahmood, Savanna A. Cohen, Gregory Corder

## Abstract

Aversion to opioid withdrawal is a significant barrier to achieving lasting opioid abstinence. The central amygdala (CeA), a key brain region for pain, threat-detection, autonomic engagement, and valence assignment, is active during opioid withdrawal. However, the role of molecularly distinct CeA neural populations in withdrawal remains underexplored. Here, we investigated the activity dynamics, brain-wide connectivity, and functional contribution of Protein Kinase C-delta (PKCδ)-expressing neurons in the CeA lateral capsule (CeLC^PKCδ^) during fentanyl withdrawal in mice. Mapping activity-dependent gene expression in CeLC^PKCδ^ neurons revealed a highly withdrawal-active subregion in the anterior half of the CeA. Fiber photometry calcium imaging showed that opioid-naïve CeLC^PKCδ^ neurons respond to salient noxious and startling stimuli. In fentanyl-dependent mice, naloxone-precipitated withdrawal increased spontaneous neural activity and enhanced responses to noxious stimuli. Chronic inhibition of CeLC^PKCδ^ neurons throughout fentanyl exposure, via viral overexpression of the potassium channel Kir2.1, attenuated withdrawal signs in fentanyl-dependent mice. Lastly, we identified putative opioid-sensitive inputs to CeLC^PKCδ^ neurons using rabies-mediated monosynaptic circuit tracing and color-switching tracers to map mu-opioid receptor-expressing inputs to the CeLC. Collectively, these findings suggest that the hyperactivity of CeLC^PKCδ^ neurons underlies the somatic signs of fentanyl withdrawal, offering new insights into the amygdala cell-types and circuits involved in opioid dependence.

## INTRODUCTION

Opioids remain gold-standard analgesics, but their side effects, including dependence and respiratory depression, limit their long-term clinical utility. Opioid dependence presents a major barrier to sustained abstinence, as abstinence or antagonist administration in dependent patients can trigger a highly aversive withdrawal syndrome that fuels relapse ^1,2^ Even with the risk of dependence, millions of pain patients rely on opioids to manage both acute and chronic pain ^3,4^. Additionally, a substantial portion of Opioid Use Disorder (OUD) patients experience comorbid chronic pain^5–7^. Understanding the interactions between long-term opioid use and the biological substrates of pain is therefore critical in addressing the opioid crisis.

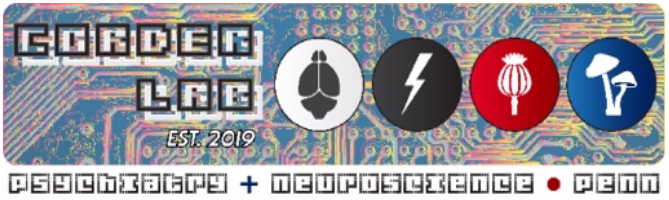

A key neural circuit within the brain pathways for pain processing, including its unpleasant, motivating aspects, is the central nucleus of the amygdala (CeA) ^8–19^. In particular, its lateral division (CeLC)—comprising the capsular nucleus (CeC) and lateral nucleus (CeL)—receives direct input from the ascending spinoparabrachial pathway, which conveys information about environmental threats and nociceptive stimuli from the periphery to the brain ^15,20,21^. CeLC neural activity coordinates diverse and rapid avoidance, escape, affective, and autonomic responses to these stimuli, helping animals avoid danger and injury ^12,22,23^. Both chronic pain and antagonist-precipitated opioid withdrawal increase neural activity markers in the CeA, including phosphorylated ERK ^24^, FOS ^14,25–27^, and calcium (Ca^2+^) activity ^28,29^. Importantly, growing evidence suggests that the CeA operates not as a single unit, but through multiple functionally, transcriptomically, and anatomically distinct cell-types with divergent responses to stimuli and often opposing effects on behavior ^14,17,18,30–40^. While recent work increasingly illustrates the diversity of CeA neurons, how these distinct cell-types may uniquely contribute to heightened CeA activity during opioid withdrawal has not been fully explored.

Protein Kinase Cδ-expressing neurons in the CeLC (CeLC^PKCδ^) have emerged as crucial regulators of aversive stimuli processing, particularly in relation to pain and threat detection ^14,15^. In rodents, CeLC^PKCδ^ neurons account for approximately half of CeLC neurons, and receive direct input from both parabrachial nucleus and valence-encoding basolateral amygdala cell-types ^34,37,41,42^. Their neuroanatomical position allows CeLC^PKCδ^ neurons to orchestrate affective and motivational responses to pain, maintaining negative emotional states in anxiety-induced anorexia and conditioned fear ^39,41,43,44^ and facilitating hypersensitivity in chronic pain models ^11,14,15,17^. Moreover, CeLC^PKCδ^ neurons exhibit heightened sensitivity to transcriptomic changes following alcohol dependence^35^, suggesting a possible role in dependence-related processes. Interestingly, a subpopulation of CeLC^PKCδ^ neurons express mu-opioid receptors (MORs), the primary molecular target of exogenous opioids such as fentanyl and morphine ^13,29^. It is unclear, however, whether CeLC^PKCδ^ neurons contribute to the various effects of opioid drugs, making them an important and compelling target for further investigation in understanding the neural circuits involved in opioid dependence and withdrawal

While the CeA is known to be involved in opioid withdrawal—due to its central role in threat detection, autonomic regulation, and negative affective state maintenance—the specific contributions of distinct neural populations, such as PKCδ-expressing neurons, is not fully understood. This study aims to fill this gap by investigating whether CeLC^PKCδ^ neurons contribute to the neural and behavioral effects of opioid withdrawal.

## RESULTS

### Precipitated fentanyl withdrawal increases cellular activity in CeLC^**PKCδ**^ **neurons**

To test whether fentanyl withdrawal increases CeLC^PKCδ^ neuron activity, we induced dependence in male and female *Prkcd*-Cre mice by providing 24-hour free access to fentanyl-treated water (0.02 mg/mL) for eight days in the homecage (**Fig. 1a**). Control mice drank untreated water (**Fig. 1a**). On the eighth day, mice received either naltrexone (1 mg/kg, s.c.) or saline. After 105 minutes, brains were collected and immunostained for FOS, an immediate early gene product used as a marker of neural activity. We found a significant effect of drinking treatment and naltrexone administration on the number of FOS^+^ nuclei in the CeA. Specifically, fentanyl-dependent mice that received naltrexone (*i*.*e*., withdrawing mice) had significantly more FOS^+^ nuclei than any other condition (*withdrawal*FOS; **Fig. 1b, c**). Fentanyl drinking alone did not significantly increase FOS^+^ nuclei compared to water-drinking/saline mice, while naltrexone alone caused a small, nonsignificant increase in FOS^+^ nuclei in water-drinking mice.

**Fig. 1.**
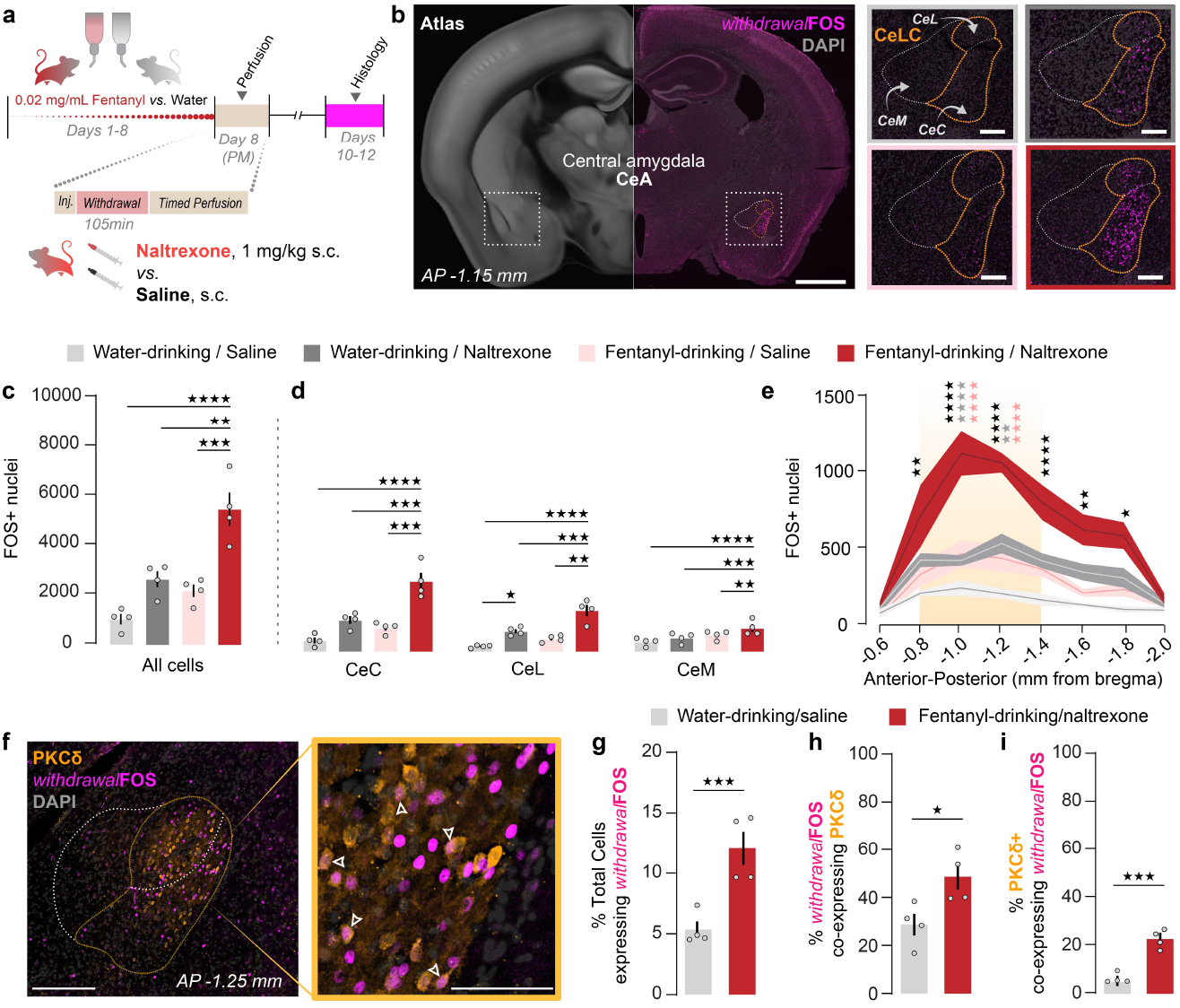
Fentanyl withdrawal increases FOS expression in CeLC^PKCδ^ neurons. **(a)** Experimental timeline. **(b)** Representative images of FOS immunoreactivity under each experimental condition. Left: Fentanyl-drinking/withdrawal, 4x-magnified. Right: 20x-magnified z-stacks. Upper left/light gray: water-drinking/saline (n=3 females, 1 male); upper right/dark grey: water-drinking/naltrexone (n=2 females, 2 males); bottom left/pink: fentanyl-drinking/saline (n=2 females 2 males); bottom right/dark red: fentanyl-drinking/naltrexone (n=3 females, 1 male). Scale bars: Left image, 1mm; right images, 250 um **(c)** Fentanyl-drinking/naltrexone mice (i.e., withdrawal mice) had significantly more FOS+ nuclei in the CeA than the mice in any other experimental condition. Two-way ANOVA with Bonferroni’s correction, significant effect of drinking condition (F(1,12)=23.80, p=0.0004) and withdrawal condition (F(1,12)=36.18, p<0.0001). Bars/error bars are mean +/-SEM; individual points represent data for a single mouse. **p<0.01; ***p<0.001; ****p<0.0001. **(d)** Fentanyl-drinking/naltrexone mice (i.e., withdrawal mice) had significantly more FOS+ nuclei in the CeC (left), the CeL (middle), and the CeM (right) subnuclei of the CeA than the mice in any other experimental condition. Water-drinking/naltrexone mice also had significantly more FOS+ nuclei only in the CeL and only compared to water-drinking/saline mice. CeC: Two-way ANOVA with Bonferroni’s correction, significant effect of drinking condition (F(1,12)=25.44, p=0.0003), withdrawal condition (F(1,12)=42.95, p<0.0001), and drinking x withdrawal interaction (F(1,12)=6.576, p=0.0248). CeL: Two-way ANOVA with Bonferroni’s correction, significant effect of drinking condition (F(1,12)=20.29, p=0.0007) and withdrawal condition (F(1,12)=47.23, p<0.0001). CeM: Two-way ANOVA with Bonferroni’s correction, significant effect of drinking condition (F(1,12)=25.03, p=0.0004), withdrawal condition (F(1,12)=18.12, p<0.0011), and drinking x withdrawal interaction (F(1,12)=12.07, p=0.0046).*p<0.05; **p<0.01; p<0.001; p<0.0001 **(e)** Fentanyl withdrawal is associated with significantly more FOS+ nuclei in the CeA than any control conditions at AP-1.0mm and AP-1.2mm from bregma. Yellow shading: approximate region targeted by viral vector injections in **Figs. 2-5**. Three-way ANOVA with Bonferroni’s correction, significant effect of coordinate (F(7,100)=17.82, p<0.0001), drinking condition (F(1,100)=60.22, p<0.0001), withdrawal condition (F(1,100)=93.90, p<0.0001) coordinate x drinking interaction (F(7,100)=3.589, p=0.0017), coordinate x withdrawal interaction (F(7,100)=3.697, p=0.0014), and drinking x withdrawal interaction (F(1,100)=10.32, p=0.0018). Asterisks: fentanyl-drinking/naltrexone vs water-drinking/saline (black); vs. water-drinking/naltrexone (grey); vs. fentanyl-drinking/water (pink). Points and area fill represent mean values +/-SEM. *p<0.05; **p<0.01; ***p<0.001; ****p<0.0001 **(f)** FOS immunoreactivity colocalizes with PKCδ immunoreactivity in the CeLC. Arrowheads: PKCδ+/FOS+ cells.. Scale bars: large image 250 um, inset 100 um. **(g)** Fentanyl withdrawal is associated with a significantly higher percentage of CeA neurons that are FOS+ (i.e., *withdrawal*FOS neurons) compared to the water-drinking/saline condition. Water-drinking/saline: n=4 females; Fentanyl-drinking/naltrexone: n=2 females and 2 males. Unpaired t-test, t=4.354, **p=0.0048. **(h)** Significantly more FOS+ neurons co-express PKCδ during withdrawal than during the water-drinking/saline condition. Unpaired t-test, t=2.877, *p=0.0282. **(i)** Significantly more PKCδ+ neurons co-express FOS during withdrawal

In withdrawing mice, *withdrawal*FOS^+^ nuclei were predominantly localized to the lateral division of the CeA (CeLC), compared to other CeA subregions, including the central lateral (CeL) and central medial (CeM) (**Fig. 1b, d**). Withdrawing mice had significantly more FOS^+^ nuclei than all other groups across all three CeA sub-nuclei (**Fig. 1d**). Notably, naltrexone in water-drinking mice selectively increased FOS^+^ nuclei in the CeL, suggesting tonic endogenous opioid activity in this region (Fig. 1d). *withdrawal*FOS expression was significantly higher in the CeLC than in the CeM across all fentanyl-drinking, naltrexone-treated mice (**Fig. S1a-b**).

*withdrawal*FOS was elevated compared to that of untreated controls throughout the anterior-posterior axis, except at the most anterior (−0.6 mm) and posterior (−2.0 mm) poles (**Fig. 1e**). However, *withdrawal*FOS expression was only significantly higher in withdrawing mice than in fentanyl-naïve, naltrexone-treated controls at −1.0 to −1.2 mm AP. Despite prior studies suggesting a greater role for the right CeA in aversive processing ^16,24,48,63^, we did not observe robust lateralization of FOS expression (**Fig. S1c– g**).

Next, we investigated whether withdrawal specifically increased FOS expression in CeLC^PKCδ^ neurons. We collected CeA tissue from untreated and withdrawing mice and co-immunostained for FOS and PKCδ (**Fig. 1f**). As expected, withdrawing mice had significantly more FOS^+^ CeLC neurons than untreated controls, with *withdrawal*FOS^+^ nuclei comprising 10–15% of total CeA nuclei in withdrawing mice compared to 5–7% in untreated controls (**Fig. 1g**). In withdrawing mice, 39–60% of FOS^+^ neurons were PKCδ+, significantly more than in untreated mice (17–38%; **Fig. 1h**). Likewise, the proportion of PKCδ ^+^ neurons that were also FOS^+^ increased significantly, from 7–11% in untreated mice to 18–26% in withdrawing mice (**Fig. 1i**).

Consistent with our initial observations, we found that the increase in FOS expression was higher in the CeC and CeL for every fentanyl-drinking, naltrexone-treated mouse compared to the CeM (**Fig. S1a**). This effect was significant when data from the entire CeLC were grouped together (**Fig. S1b**).

Evidence suggests that the right hemisphere CeA plays a greater role in nociceptive processes, contributing more to the generation of pain-related aversive behaviors and responses to nociceptive stimuli than the left ^16,24,48,63^. We hypothesized that aversive opioid withdrawal would also result in increased withdrawal-induced FOS expression in the right CeA. A repeated measures two-way ANOVA revealed a significant effect of hemisphere when analyzing the CeA as a whole, but multiplicity-adjusted post hoc tests did not show any significant differences between hemispheres under any treatment condition (**Fig. S1c**). We did not find a main effect of hemisphere in the CeC (**Fig. S1d**), but we did find a main effect of hemisphere in the CeL and CeM. However, only in the water-drinking/naltrexone group (in the CeL; **Fig. S1e**) and the fentanyl-drinking/naltrexone group (**Fig. S1f**) were these effects significant after multiplicity-adjusted posthoc tests. When the CeLC was considered as a whole, we did not detect a main effect of hemisphere (**Fig. S1g)**. Therefore, while there were subtle differences in the left and right side of the CeA in terms of *withdrawal*FOS expression, we did not detect a robust lateralization effect, such as those reported previously in the context of pain ^16,24,48,63^.

### CeLC^PKCδ^ neurons are tuned to salient aversive and noxious stimuli

Based on our initial FOS mapping, which indicated increased neural activity from −1.0 to −1.2mm AP during fentanyl withdrawal (**Fig. 1e**), we next investigated the temporal dynamics of anterior CeLC^PKCδ^ neurons using in vivo fiber photometry in behaving mice. We expressed the Ca^2+^ indicator GCaMP6f in this region of the CeA of *Prkcd*-Cre mice and placed an optical fiber 0.2 mm above the injection site to measure stimulus-evoked fluorescence changes (**Fig. 2a, b**). Although several studies demonstrate the role of CeLC^PKCδ^ neurons in generating and responding to aversive states ^14,17,37,44,64^, few have directly examined their immediate responses to discrete noxious and aversive stimuli.

**Fig. 2.**
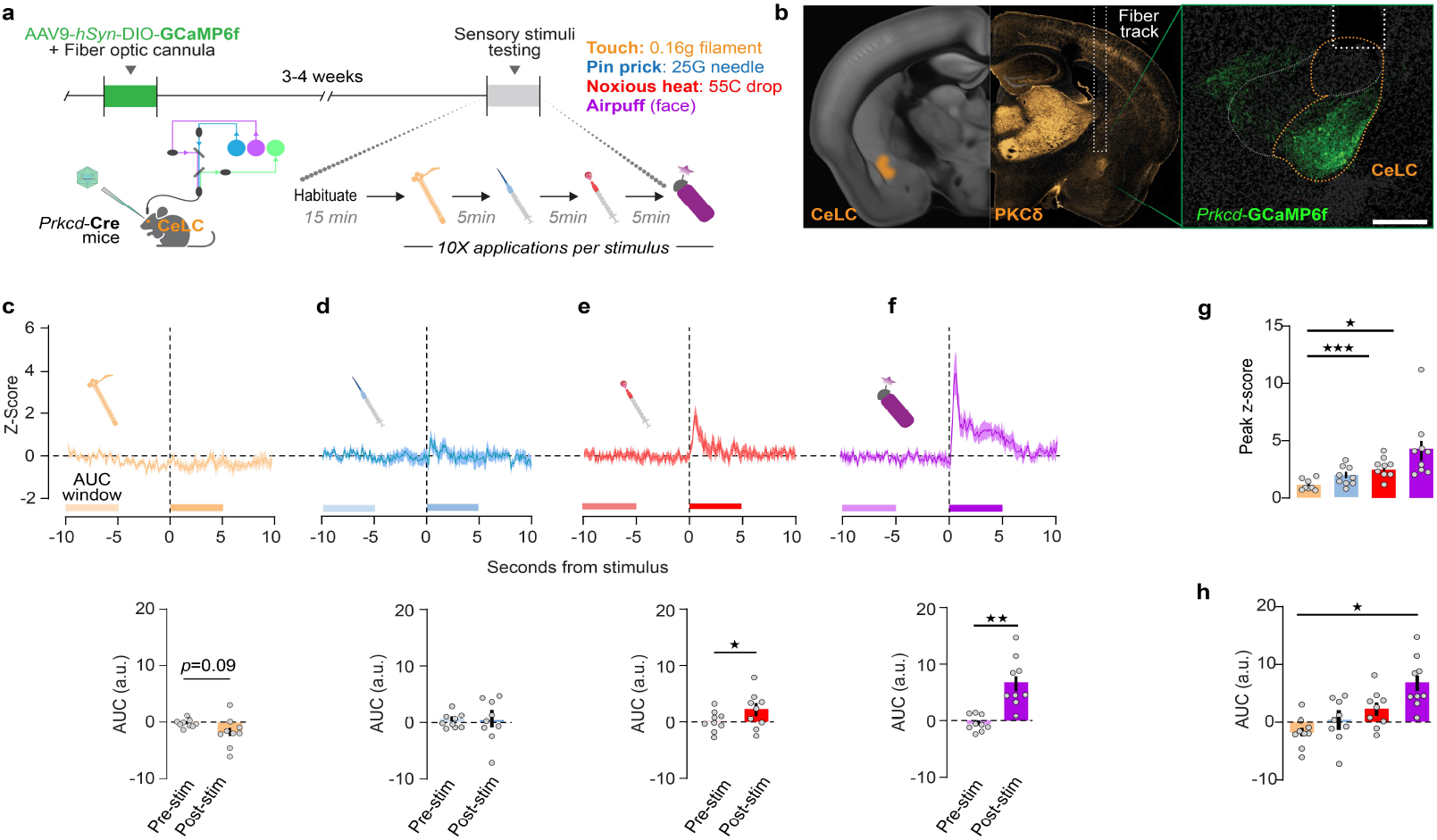
CeLC^PKC^^δ^ neurons in opioid-naïve mice respond to noxious aversive and non-noxious aversive stimuli. **(a)** Experimental timeline. **(b)** Left: PKCδ immunoreactivity and approximate location of fiber optic cannula in the targeted area of the right hemisphere. Right: expression of AAV9-hsyn-DIO-GCaMP6f in the CeLC of a *Prkcd*-Cre mouse, with a fiber optic positioned approximately 200 um dorsal to the injection site. Scale bar: 250 um. **(c)-(f)** Top: Peri-stimulus time histogram of normalized dF/F (z-score) from −10 sec to 10 sec from the moment of stimulus application, following application of **(c)** a 0.16 g von Frey filament (innocuous light touch); **(d)** pinprick with a 25-gauge needle (noxious pinprick); **(e)** a 55°C hot water drop (noxious hot water); and **(f)** an aversive, but non-noxious, airpuff delivered to the side of the face contralateral to virus injection and fiber (aversive airpuff). Bottom: Area under the z-score curve from 10-5 seconds prior to stimulus application vs. from 0-5seconds after stimulus application. **(c)** Compared to pre-stimulus baseline, innocuous light touch was associated with a small, nonsignificant decrease in the AUC (Paired t-test, t=1.902). Bars represent mean z-score +/-SEM; dots represent individual points. **(d)** Noxious pinprick did not significantly affect AUC. **(e)** Noxious hot water was associated with a significant increase in AUC (Unpaired t-test, t=2.697, *p=0.0412). **(f)** Aversive airpuff was associated with a significant increase in AUC (Unpaired t-test, t=4.9825, \*\**p*=0.013). Lines and area fill represent mean values of 10 trials/subject, averaged across all subjects, +/-SEM (n=4 females and 5 males). **(g)** Noxious hot water and aversive non-noxious airpuff produced a significantly higher peak than innocuous light touch (One-way repeated measures ANOVA with Bonferroni’s correction, significant effect of treatment (F(1.544, 12.35)=20.43, p=0.0002) and significant effect of subject (F(8,24)=9.072, p<0.0001) and airpuff produced an increased peak, but this did not reach statistical significance. \**p*=0.0405; \*\*\**p*=0.001 **(h)** Compared to innocuous light touch, aversive airpuff produced a significant increase in the AUC (Mixed-effects analysis with repeated

To address this, we measured Ca^2+^ responses to four somatosensory stimuli: innocuous light touch (0.16g von Frey filament), noxious, aversive stimuli (25-gauge sharp pinprick or 55°C hot water drop), and a non-noxious but aversive stimulus (1-sec airpuff; **Fig. 2c-f**). Innocuous light touch resulted in a small, non-significant decrease in z-scored Ca^2+^ signal, dropping below baseline ∼5 sec before the stimulus and remaining suppressed for ∼8 sec post-stimulus (**Fig. 2c**). In contrast, all aversive stimuli produced a Ca^2+^ increase (**Fig. 2d-f**). The signal following noxious pinprick and hot water returned to baseline within ∼3 sec (**Fig. 2d, e**), whereas the response to aversive airpuff remained elevated for ∼8 sec post-stimulus (**Fig. 2f**). Compared to innocuous light touch, a noxious hot water drop and an aversive but non-noxious airpuff produced a significantly higher peak z-scored Ca^2+^ response in the 5 sec following stimulus application. (**Fig. 2g**). Non-normalized dF/F followed the same pattern (**Fig. S2b–f**). We also analyzed the area under the z-scored Ca^2+^ curves (AUC) before and after each stimulus. AUC slightly decreased pre- and post-innocuous light touch (**Fig. 2c**), while no significant changes were observed for noxious pinprick (**Fig. 2d**). Both noxious hot water (**Fig. 2e**) and airpuff (**Fig. 2f**) significantly increased AUC relative to baseline, with the AUC post-aversive airpuff also significantly greater than post-innocuous light touch (**Fig. 2h**). Non-normalized dF/F followed similar patterns, but only aversive airpuff produced a significant post-stimulus increase or differed significantly from innocuous light touch (**Fig. S2g–k**). From these findings, we conclude that in opioid-naïve mice, CeLC^PKCδ^ neurons primarily respond to aversive stimuli but do not differentiate noxious vs. non-noxious aversive stimuli.

### Opioid administration and precipitated withdrawal dynamically modulate CeLC^PKCδ^ neuron activity

We next examined how CeLC^PKCδ^ neurons respond to aversive stimuli during fentanyl withdrawal. (**Fig. 3a**). To render mice dependent, we used the same 24-hr access oral fentanyl- or water-drinking procedure described in **Fig. 1a**. On the eighth day of fentanyl consumption, all mice received naloxone (3 mg/kg, s.c.). As expected, fentanyl-drinking mice exhibited significantly higher global withdrawal scores than water-drinking mice following naloxone (**Fig. 3b**). In opioid-naïve mice, Ca^2+^ activity transiently dropped below baseline following naloxone injection before returning to pre-drug levels within ∼10 minutes (**Fig. 3c-d, Fig. S3a**). However, in fentanyl-dependent mice, Ca^2+^ activity did not decrease but instead rose sharply during and after naloxone administration, remaining significantly elevated throughout the 15-minute recording period (**Fig. 3f, Fig. S3b**). These findings build on our previous results (**Fig. 1**), demonstrating a rapid and sustained increase in CeLC^PKCδ^ neuron activity during precipitated opioid withdrawal.

**Fig. 3.**
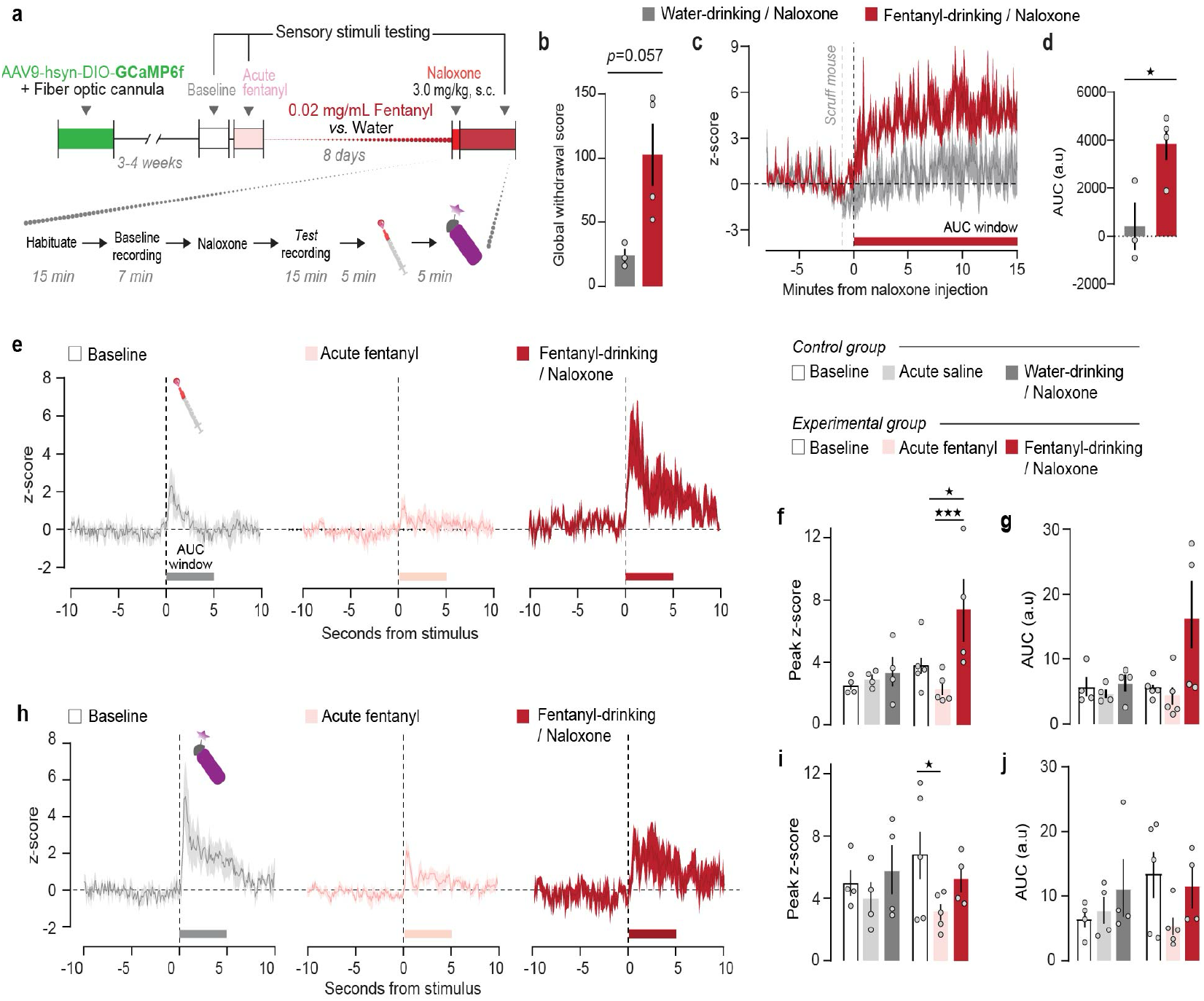
CeLC^PKC^^δ^ neurons are hyperactive and hypersensitive during fentanyl withdrawal. **(a)** Experiment timeline. After baseline sensory testing **(Fig. 3)**, mice were separated into a control group that remained opioid-naïve prior to naloxone administration on day 8 (control group; n=3 females and 1 male), and an experimental group that received all fentanyl treatments (experimental group; n=2 females and 3 males). **(b)** Fentanyl-drinking mice exhibited a trend towards higher global withdrawal scores following naloxone administration compared to water-drinking control mice.(Mann-Whitney test) **(e)** Peri-injection time histogram of fluorescence before, during, and after an injection of 3 mg/kg naloxone in control, naïve animals (dark gray) and fentanyl-dependent animals (red). Red bar: AUC interval in **(d)**. n=3 (control group; 1 male excluded) and n=4 (experimental group; 1 female excluded). **(d)** The net AUC for the 15min after naloxone injection is significantly higher for fentanyl-drinking vs. naïve mice (Unpaired t-test, t=3.008, *p=0.0298). **(e)** Peri-stimulus time histogram from 10 seconds prior to 10seconds following application of a noxious hot water drop for the fentanyl-dependent group at baseline (light grey trace), after 0.2 mg/kg fentanyl prior to drinking (pink), and during fentanyl withdrawal (red). Lines and area fill represent mean values of five trials averaged across subjects. **(f)** The peak of the fluorescence response in the 5 seconds following application of noxious hot water was significantly higher during fentanyl withdrawal than at baseline or after an acute injection of fentanyl in the experimental group (Mixed-effects analysis with Bonferroni’s adjustment, main effect of experimental timepoint (F(2,13) = 7.268, p=0.0076) and treatment x timepoint interaction (F(2,13)=4.704, p=0.029) *p=0.0133, **p=0.001 **(g)** No significant increases in 0-5 second post-stimulus AUC during any experimental condition in either group. **(h)** Peri-stimulus time histogram from 10 seconds prior to 10 seconds following application of a noxious hot water drop for the fentanyl-dependent group. **(i)** Acute fentanyl injection decreased the peak of the fluorescence response to airpuff relative to baseline in the experimental group (Mixed-effects analysis with Bonferroni’s adjustment, main effect of experimental timepoint (F(2, 13)=4.454, p=0.0336) *p=0.0215 **(j)** No significant changes in 5 second AUC during any experimental condition in either group.

To determine how fentanyl alters CeLC^PKCδ^ responses to aversive stimuli, we measured Ca^2+^ responses to a noxious hot water drop (**Fig. 3e–g**) and an aversive airpuff (**Fig. 3h–j**) under baseline conditions, after an acute fentanyl injection (0.2 mg/kg, s.c.) while mice were opioid-naïve, and during precipitated withdrawal. An experimental group received both the acute injection of fentanyl and fentanyl-treated water, while a control group received s.c. saline instead of 0.2 mg/kg fentanyl and drank untreated water – thus remaining opioid-naïve prior to the administration of naloxone on drinking day 8. Following the hot water drop, peak z-scored Ca^2+^ responses were significantly higher during fentanyl withdrawal compared to both baseline and acute fentanyl conditions (**Fig. 4e, f**). In contrast, acute saline or naloxone administration in opioid-naïve animals did not significantly alter the response (**Fig. 4e, f; S3c, d**). We also observed a consistent but nonsignificant decrease in peak Ca^2+^ responses following 0.2 mg/kg fentanyl, with no effect of saline or naltrexone in water-drinking mice. A similar trend was seen in the area under the curve (AUC) during the first 5 seconds post-stimulus, though the main effect of treatment timepoint was not significant (**Fig. 4g, Fig. S3e**). For the aversive airpuff, peak z-scored Ca^2+^ responses were significantly lower following acute fentanyl administration but remained unchanged during withdrawal (**Fig. 4h, I, Fig. S3f)**. AUC analyses showed no significant differences in either fentanyl-treated or water-treated mice (**Fig. 4j**). Non-normalized dF/F values followed similar patterns but lacked statistical significance (**Fig. S3g, h**). Thus, our results indicate that CeLC^PKCδ^ neurons exhibit heightened sensitivity to noxious stimuli during fentanyl withdrawal but reduced responsiveness to an airpuff following an analgesic dose of fentanyl.

**Fig. 4.**
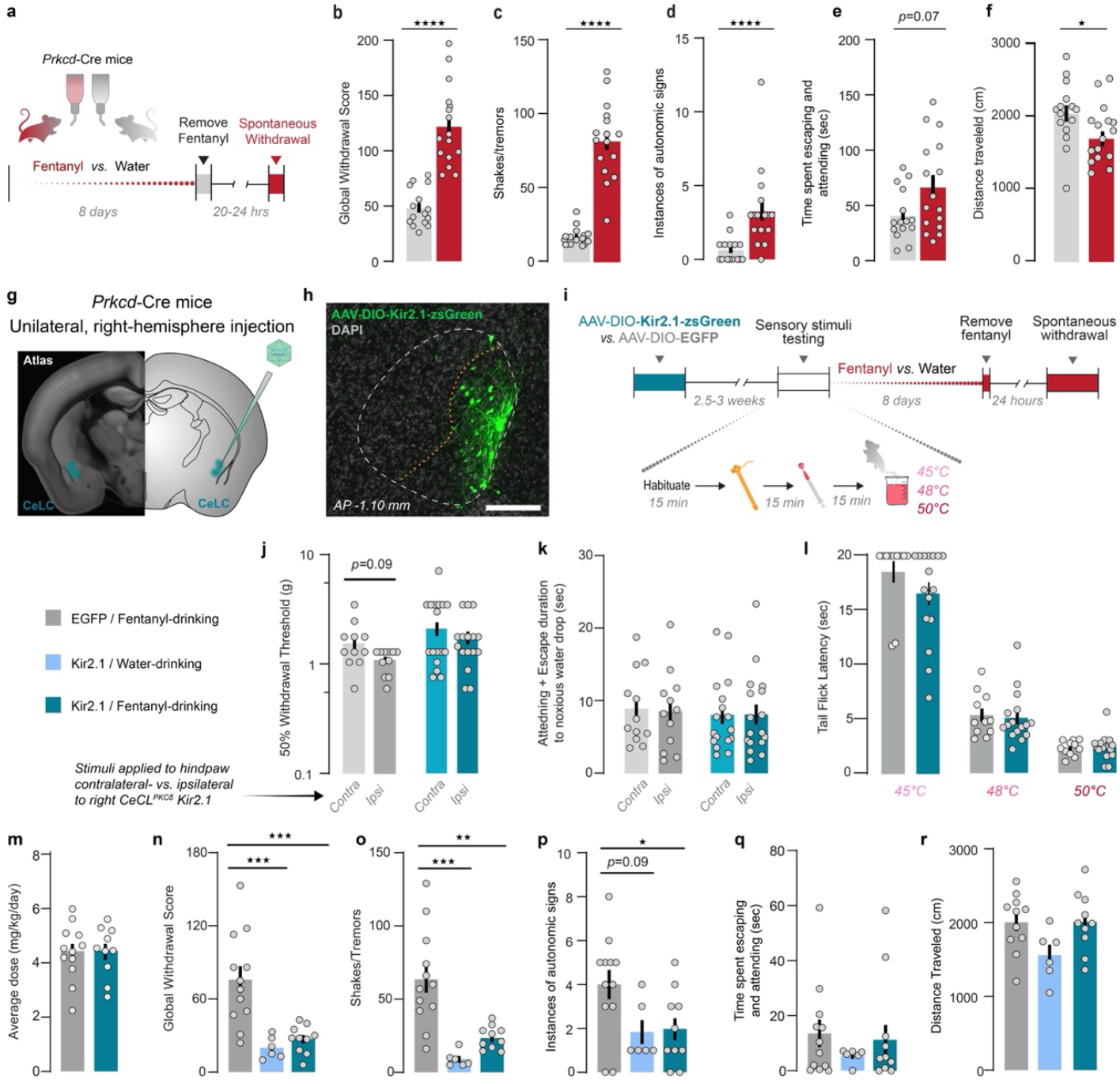
Chronic inhibition of CeLC^PKCδ^neurons prevents fentanyl withdrawal without impacting pain sensitivity. **(a-f)** Characterization of spontaneous withdrawal behavioral phenotype in *Prkcd-*Cre mice. **(a)** Experimental timeline. **(b)** Fentanyl-drinking mice exhibited significantly higher global withdrawal scores 20-24 hrs after the removal of fentanyl compared to water-drinking mice. Unpaired t-test, t=7.916, ****p<0.0001. Bars represent mean +/-SEM; dots are individual subjects’ data points. Fentanyl-drinking mice exhibited significantly more **(c)** shakes and tremors (Unpaired t-test, ****p<0.0001) and (**d**) autonomic signs (Mann-Whitney test, U=27.50, ****p<0.0001). **(e)** Fentanyl-drinking mice did not exhibit significantly higher time engaging in affective responses during withdrawal compared to water-drinking mice (Mann-Whitney test). **(f)** Fentanyl-drinking mice traveled a significantly lower distance than water-drinking mice during the spontaneous withdrawal observation period (Unpaired t-test, 2.293, *p=0.0293). **(g)-(r)** Effects of Kir2.1 overexpression in CeLC^PKCδ^ neurons. **(g)** Experimental approach. **(h)** Expression of AAVDJ-CMV-DIO-Kir2.1-zsGreen in the CeLC of *Prkcd-*Cre mice. Scale bar: 250 um. **(i)** Experimental timeline and testing procedure for sensory testing. **(j)** We detected a main effect of stimulation side (Two-way ANOVA with Bonferroni’s correction, main effect of stimulation side (F(1, 25)=7.274, p=0.0123), effect of subject (F(25,25)=3.721, p=0.0008) but not viral condition on 50% withdrawal thresholds in the von Frey Up-Down assay. Mice had slightly (but not significantly) higher 50% withdrawal thresholds on the hindpaw contralateral to the hemisphere of virus injection compared to the ipsilateral paw, but this was true only for EGFP-expressing mice. Grey bars: EGFP-expressing mice; Blue/teal bars: Kir2.1-expressing mice. Bars represent mean values +/-SEM; grey points represent individual subjects’ values. EGFP: n=6 females and 6 males. Kir2.1: n=9 females and 7 males. (**k)** Kir2.1 overexpression does not affect the time spent engaging in affective responses to the application of a noxious hot water drop, regardless of which hindpaw is stimulated (Two-Way repeated measures ANOVA). **(l)** Kir2.1 overexpression does not impact the latency for mice to withdraw tails from hot water at 3 different temperatures (Two-Way Repeated Measures ANOVA with Bonferroni’s correction, main effect of temperature (F(2,54)=165.9; p<0.0001) **(m)** Kir2.1-overexpressing mice consumed a similar average daily dose of fentanyl as EGFP-expressing control mice (Unpaired t-test). EGFP + Fentanyl-drinking: n=6 females and 6 males. Kir2.1 + Fentanyl-drinking: n=6 females and 4 males. **(n)** Kir2.1-overexpressing mice (teal) show significant fewer withdrawal signs after fentanyl exposure compared to EGFP-expressing mice (grey), and similar global withdrawal scores to Kir2.1-expressing mice that drank water (light blue; One-way ANOVA with Bonferroni’s correction, F(2,25) = 6.116), p=0.0069; ***p<0.001). EGFP + Fentanyl-drinking: n=6 females and 6 males. Kir2.1 + Fentanyl-drinking: n=6 females and 4 males. Kir2.1 + water-drinking: n=3 females and 3 males. **(o)** Kir2.1-expressing, fentanyl-drinking mice and Kir2.1-expressing, water-drinking mice exhibited significantly fewer shakes and tremors compared to EGFP-expressing, fentanyl-drinking mice (One-way ANOVA with Bonferroni’s correction, F(2,25)=5.854, p<0.0001). **p=0.001, ***p=0.0002. **(p)** Kir2.1 expression significantly reduced the autonomic subscore of fentanyl-dependent mice (One-way ANOVA with Bonferroni’s correction, (F(2,27)=5.083, p=0.0134; *p=0.01) **(q)** Kir2.1 expression did not significantly reduce the time spent engaging in affective behaviors in fentanyl-dependent mice (One-way ANOVA). **(r)** No significant effect of Kir2.1 overexpression on withdrawal-associated hypolocomotion (One-way ANOVA, F(2,24)=3.088, p=0.064).

### Behavior profile of spontaneous withdrawal from fenta-nyl drinking

We next sought to understand whether the increase in neural activity seen during fentanyl withdrawal was necessary for dependence. To do so, we first characterized the spontaneous withdrawal phenotype of our fentanyl drinking procedure in *Prkcd*-Cre mice 20–24 hours after the removal of the fentanyl-treated water (**Fig. 1a, Fig. 4a**). Fentanyl-drinking mice consumed a similar volume of water over the 8-day period as water-drinking mice (**Fig. S4a**). Neither male nor female mice experienced significant weight loss during the 8-day administration paradigm, indicating that they received sufficient hydration (**Fig. S4b**). While doing so, fentanyl-drinking mice consumed an average of 3.89 mg/kg/day (**Fig. S4c**). There was a significant effect of Treatment Day and subject on intake volume, with both water- and fentanyl-drinking mice consuming a significantly higher volume on Treatment Day 7, and fentanyl-drinking mice consuming significantly more on Treatment Day 8, compared to day 1 (**Fig. S4c-d**). There was a general upward trend in consumed dose, and on Treatment Day 8 all mice consumed more fentanyl than on Treatment Day 1 (**Fig.S4e**).These effects were present in both female and male mice, with a significant sex x treatment day interaction but no main effect of sex, with female mice consuming a slightly higher dose of fentanyl compared to male mice (**Fig. S4f**). However, we noticed similar patterns of drinking in water-drinking mice, suggesting that this result was due to the experimental conditions rather than representing volitional dose escalation. Notably, this drinking procedure did not produce a significant preference for fentanyl-treated vs. untreated water in a two-bottle choice procedure: on average, mice consumed similar quantities from both bottles throughout treatment (**Fig. S4g-h**).

On Treatment Day 8, the fentanyl-treated water was replaced with untreated water for fentanyl-drinking mice. We then assessed spontaneous withdrawal 20-24 hrs later, on Day 9. *Prkcd*-Cre mice exhibited typical signs of opioid withdrawal, with fenta-nyl-drinking mice displaying significantly higher global withdrawal scores than water-drinking mice (**Fig. 4b**). This effect was present in both sexes, and we did not detect a significant effect of sex on global withdrawal scores (**Fig. S4f**). To determine which withdrawal signs drive the global withdrawal scores in our model, we calculated subscores based on three broad categories of typical opioid withdrawal, present in both people and rodent models: shakes and tremors, autonomic symptoms, and affective symptoms (**Fig. 4c-e**). In our model, fentanyl withdrawal was characterized by a significant increase in the number of shakes and tremors (**Fig. 4c**), including significant increases in resting tremor, body shakes, head shakes, and paw tremors (**Supplementary Fig. S4k-n**). In addition, fentanyl-drinking mice exhibited more autonomic withdrawal symptoms than water-drinking mice (**Fig. 4d**): they excreted significantly more feces (**Fig. S4o**) and exhibited more frequent teeth chattering (**Fig. S4p**). Overall, fentanyl-drinking mice did not exhibit significantly more affective behaviors than water-drinking mice (**Fig. 4e**). They did spend more time in genital grooming, a frequently-reported withdrawal-associated behavior (**Fig. S4q**) but we did not detect significantly more jumping (**Fig. S4r**), defensive treading (**Fig. S4s**), digging (**Fig. S4t**), or limb biting (**Fig. S4u**). Fentanyl-drinking mice also displayed significantly lower locomotor activity during the 15min spontaneous withdrawal period (**Fig. 4f**). Thus, our fentanyl-drinking paradigm is sufficient to produce dependence in male and female *Prkcd*-Cre mice, indicated by the development of the spontaneous withdrawal syndrome after the removal of fentanyl.

Chronic opioid treatment can produce paradoxical hyperalgesia^65,66^. In addition, new or returning pain can accompany opioid withdrawal in clinical populations ^66,67^. We therefore assessed nociceptive sensory-reflexive thresholds and affective-motivational behaviors in response to mechanical and thermal stimuli, both on Treatment Day 6 of fentanyl and immediately after the spontaneous fentanyl withdrawal assessment (**Fig. S5a**). We did not detect an effect of treatment condition or experimental timepoint on 50% withdrawal thresholds in the von Frey Up-Down assay (**Fig. S5b**). While we detected a significant effect of experimental timepoint on tail withdrawal latency from 48°C and 50°C water, wherein a gradual decline in withdrawal latencies occurred over the 3 Treatment Days, multiplicity-adjusted tests did not detect any significant differences in fentanyl-treated mice between timepoints for either water temperature (**Fig. S5c**). We also did not see an effect on the time mice engaged in affective responses to a hot water drop applied to the left hindpaw (**Fig. S5d**) or an inescapable hotplate (**Fig. S5e**). Thus, our fentanyl-drinking procedure does not produce robust fentanyl-induced hyperalgesia or withdrawal-induced hyperalgesia.

### CeLC^PKCδ^ Neuron Inhibition Does Not Alter Pain Sensitivity but Prevents Fentanyl Withdrawal Symptoms

To assess whether CeLC^PKCδ^ neuron activity is required for fentanyl dependence, we chronically inhibited these neurons using the potassium channel Kir2.1. We injected *a* Cre-dependent Kir2.1-zsGreen (AAVDJ-*CMV*-DIO-Kir2.1-zsGreen) or a control EGFP-expressing vector (AAV5-*hSyn*-DIO-EGFP) into the CeLC of *Prkcd-*Cre mice before fentanyl exposure (**Fig. 4g-i, Fig. S6a**). Given prior evidence that CeLC^PKCδ^ neurons contribute to nociceptive processing^13,14,68^ and our fiber photometry results showing that these neurons respond to hot water application in opioid-naïve mice, we first tested whether Kir2.1 overexpression affected pain sensitivity. There were no significant differences in mechanical (**Fig. 4j**) or thermal (**Fig. 4l**) withdrawal thresholds between EGFP- and Kir2.1-expressing mice, nor any differences in von Frey thresholds between the contralateral and ipsilateral paws relative to the virus injection (**Fig. 4i**). Additionally, Kir2.1 overexpression did not significantly alter escape behavior or attentive responses to a 55°C hot water drop on either hindpaw (**Fig. 4k**). Consistent with previous observations that inhibition of CeLC^PKCδ^ neurons impacts nociceptive responses only after injury ^14^, these findings indicate that CeLC^PKCδ^ neurons baseline activity is not required for mechanical or thermal nociception in opioid-naïve mice.

### CeLC^PKCδ^ Neuron Inhibition Prevents Spontaneous Fen-tanyl Withdrawal Symptoms

Next, we assessed whether Kir2.1-expressing mice developed spontaneous fentanyl withdrawal. Importantly, both EGFP- and Kir2.1-expressing mice consumed comparable amounts of fentanyl (**Fig. 4m**). Kir2.1 expression prevented withdrawal symptoms, as fentanyl-drinking, Kir2.1-expressing mice did not show significantly higher withdrawal scores than water-drinking controls, and their scores were significantly lower than those of EGFP-expressing, fentanyl-drinking mice (**Fig. 4n**). As expected, EGFP-expressing, fentanyl-drinking mice exhibited significantly higher global withdrawal scores than water-drinking, Kir2.1-expressing mice. This pattern was also observed for shakes/tremors (**Fig. 4o**) and autonomic withdrawal symptoms (**Fig. 4p**), though this effect was not statistically significant for the autonomic subscore. Additionally, affective withdrawal symptoms did not significantly differ between groups (**Fig. 4q**). The primary withdrawal-related behaviors that were reduced by Kir2.1 expression were resting tremors and paw tremors (**Fig. S6b–l**). Lastly, while fentanyl withdrawal typically reduces locomotor activity (**Fig. 4f**), we observed a small but nonsignificant increase in locomotion in both fentanyl-drinking groups compared to water-drinking, Kir2.1-expressing mice (**Fig. 4r**). This suggests that CeLC^PKCδ^ inhibition may modulate locomotor activity in the absence of fentanyl withdrawal.

### Brain-wide identification of opioid-sensitive inputs to CeLC^**PKCδ**^ **neurons**

Our findings indicate that CeLC^PKCδ^ neurons respond to both pre-cipitated and spontaneous withdrawal from fentanyl. To determine whether these effects are mediated by opioid-sensitive inputs to CeLC^PKCδ^ neurons, we used viral-mediated circuit tracing to identify brain regions that meet two criteria: (1) provide monosynaptic input to CeLC^PKCδ^ neurons, and (2) express MORs on CeA-projecting neurons.

To map monosynaptic inputs to CeLC^PKCδ^ neurons, we performed rabies-mediated retrograde tracing in *Prkcd*-Cre mice (**Fig. 5a– c**). We injected a mixture of Cre-dependent TVA-mCherry and G-protein “helper” viruses into the the CeLC of *Prkcd-*Cre mice 2 weeks prior to the injection of EnvA-pseudotyped RABV*d*G-GFP, which then selectively enters PKCδ+ neurons via the TVA receptor and retrogradely “hops” only one synapse using the G-protein.

**Fig. 5.**
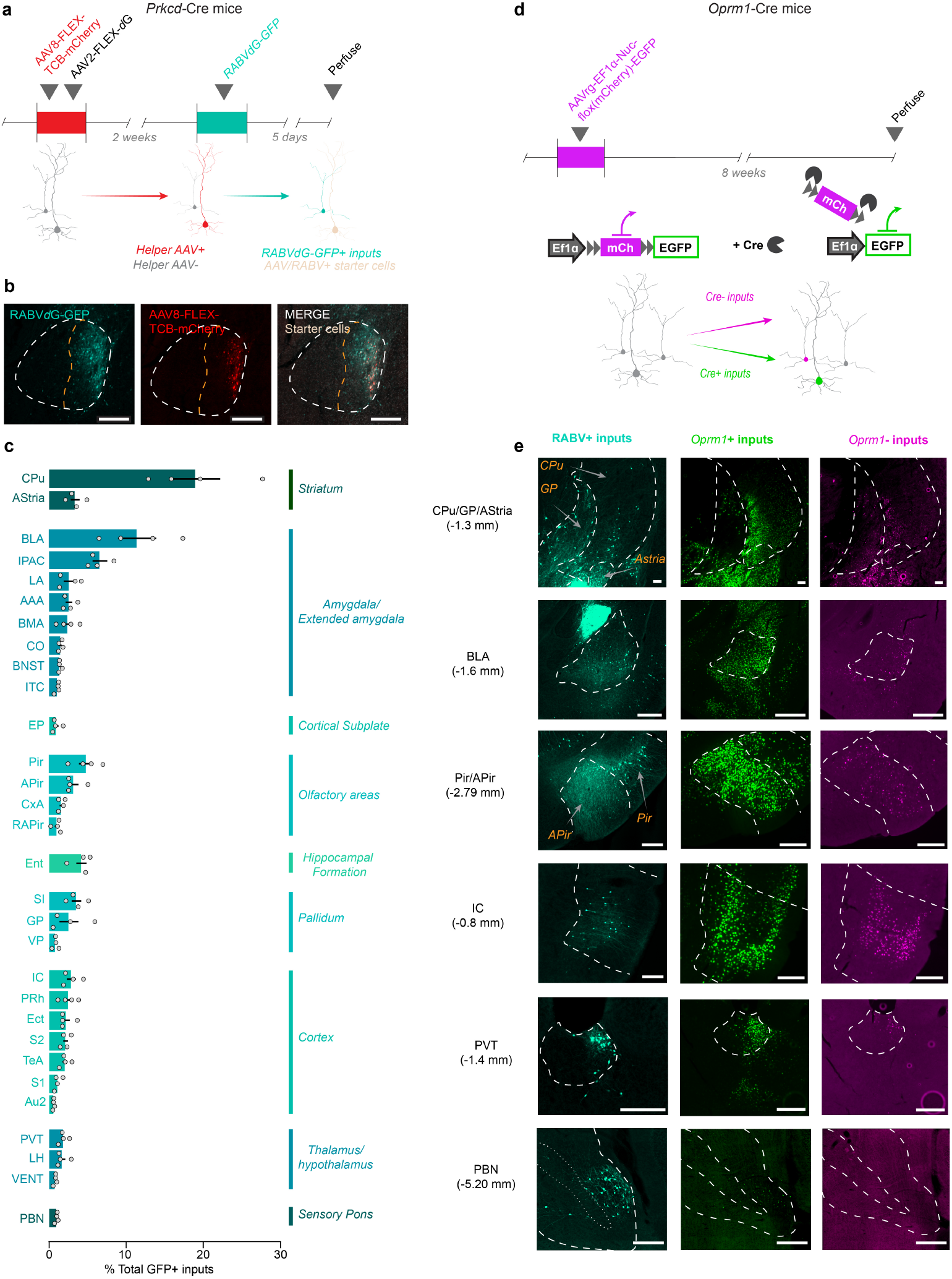
Presynaptic input architecture of opioid-receptor-expressing circuits to CeLC^PKCδ^ neurons. **(a)** Timeline for the expression of RABV*d*G in the CeA of *Prkcd*-Cre mice. Cre+ cells are initially transfected with Cre-dependent “helper” AAVs containing a modified TVA receptor tagged with mCherry, and the rabies glycoprotein. When RABV*d*G-GFP is injected into the CeA 2 weeks later, it can only transfect cells that took up the helper AAVs (“starter cells”; tan). RABV*d*G-GFP moves one synapse back, labeling direct inputs to the “starter cells” with GFP (cyan). **(b)** Expression of RABV*d*G-GFP and the fluorescently-labeled helper virus AAV8-hsyn-FLEX-TCB-mCherry containing the modified TVA CeLC^PKCδ^ in the CeA. Scale bars: 250u m. **(c)** Top 30 regions identified with the highest densities of inputs to CeLC^PKCδ^ neurons (i.e., RABV*d*G-GFP-labeled cells). Bars represent mean percentage of identified GFP+ cells +/-SEM; dots represent values for individual subjects. n=2 females and 2 males **(d)** Experimental timeline for the expression of AAVrg-EF1*α*-Nuc-flox(mCherry)-EGFP in *Oprm1*-Cre mice. In the presence of Cre, mCherry will be deleted due to the orientation of loxP sites; *Oprm1*-Cre+ cells thus express the EGFP. In the absence of Cre, mCherry is not deleted, and a stop codon downstream of the mCherry prevents the expression of EGFP; *Oprm1*-Cre-cells thus express mCherry. **(e)** Representative images of RABV*d*G-GFP-labeled (cyan), GFP-labeled Cre+ cells (green), and mCherry-labeled Cre-cells (magenta) in notable input regions. Scale bars: 200 um. Abbreviations: CPu, caudate-putamen; AStria, amygdalostriatal transition area; BLA: basolateral amygdal; IPAC: Interstitial nucleus of the anterior limb of the posterior commissure; LA: lateral amygdala; AAA: anterior amygdaloid area; BMA: basomedial amygdala; CO: cortical amygdala; BNST: Bed nucleus of the stria terminalis; ITC: intercalated nuclei of the amygdala; EP: endopiriform nucleus; Pir: Piriform cortex; APir: amygdalopiriform transition area; CxA: cortex-amygdala transition area; RAPir: rostral amygdalopiriform area; Ent: Entorhinal cortex; SI: Substantia innominata; GP: Globus pallidus, VP: ventral pallidum; IC: insular cortex; PRh: Perirhinal cortex; Ect: Ectorhinal cortex; S2: Secondary somatosensory cortex; TeA: Temporal association cortex; S1: Primary somatosensory cortex; AuV/AuD: Ventral and Dorsal auditory cortex; PVT: Paraventricular nucleus of the thalamus; LH: Lateral hypothalamus: VENT: Ventral posterior thalamic nuclear group; PBN: Parabrachial nucleus.

Consistent with previous studies^38^, we observed dense monosynaptic inputs from the striatum, amygdala, and extended amygdala, as well as substantial projections from sensory cortical areas, including olfactory, somatosensory, and auditory regions (**Fig. 5c**). We also identified inputs from the thalamus, hypo-thalamus, and pallidum, with the parabrachial nucleus (PBN) serving as the primary hindbrain input (**Fig. 5c**). Several of these regions, including the insular cortex^69–71^, dorsal striatum^72^, basolateral amygdala^73,74^, and paraventricular thalamus ^75,76^, are well-known mediators of drug dependence. Notably, we did not detect labeled cells in the contralateral CeA, suggesting minimal interhe-mispheric connectivity within the CeLC^PKCδ^ population.

Next, we used a retrogradely trafficked, Cre-dependent color-switching viral tracer (AAVretrograde-*EF1α*-Nuc(flox)-mCherry-EGFP) in *Oprm1*-Cre mice to assess the density of MOR-expressing inputs to the CeA in the regions identified via RABV*d*G-mediated tracing. This approach labels the nuclei of *Oprm1*^+^ inputs with EGFP and Oprm1− inputs with mCherry (**Fig. 5d**). The distribution of inputs in both hemispheres was consistent between RABVdG and color-switching tracer mice (**Fig. 5e, Fig. S7, Fig. S8**). All examined brain regions contained both *Oprm1*^+^ and *Oprm1*^−^ inputs. Some areas, such as the insular cortex, basolateral amygdala, and ventral posterior thalamus, had a greater proportion of *Oprm1*^−^ inputs, but still contained *Oprm1*^+^ projections (**Fig. 5e, Fig. S7**). Both tracing strategies labeled comparatively few contralateral inputs to the CeA (**Fig. S8**)

### Parabrachial MOR-expressing neurons project directly to CeLC^**PKCδ**^ **neurons**

CeLC^PKCδ^ neurons are the ascending target of the spinoparabrachial pathway, and the parabrachial nucleus→CeLC input has a well-established role in processing and responding to noxious and non-noxious aversive stimuli.^15,36,48,77^ Additionally, the parabrachial nucleus, and particularly its external lateral subnucleus, expresses MORs at high density (**Fig. S9a**). Our viral tracing indicates that *Oprm1*^+^ neurons in the PBN project to the CeA, but it is unclear whether *Oprm1*^+^ neurons synapse on CeLC^PKCδ^ neurons. To address this, we more thoroughly characterized the inputs from the parabrachial nucleus to the CeA and to CeLC^PKCδ^ neurons in particular. Many subnuclei of the PBN express MORs, but they are particularly densely expressed in the external lateral region of the PBN (elPBN; **Fig. S9a**). As expected ^21^, the injection of a virus containing a Cre-dependent fluorophore into the PBN of *Oprm1*-Cre mice results in terminal labeling in the CeLC, including in the anterior portion of the CeC where *withdrawal*FOS+ neurons are particularly dense (**Fig. S9b-d**). Our RABV*d*G tracing also revealed monosynaptic inputs to CeLC^PKCδ^ neurons that were primarily located in the elPBN (**Fig. S9e**). To determine whether the MOR+ terminals might synapse on CeLC^PKCδ^ neurons, we coupled RABV*d*G-GFP tracing with *in situ* hybridization to label *Oprm1* mRNA in RABV*d*G-GFP-labeled cells in the PBN (**Fig. S9f-g**). We determined that the majority (42/72 identified cells, roughly 58%) of PBN monosynaptic inputs to CeLC^PKCδ^ neurons express the mu opioid receptor (**Fig. S9g**).

## DISCUSSION

Neural activity within the CeA is crucial for processing negative emotional states and aversive experiences, including those linked to opioid withdrawal. Though recent research increasingly illuminates the diversity of CeA cell-types and their associated neural circuits, significant gaps remain regarding how the specific cell-types and pathways in the CeA sustain opioid dependence. Using cell-type-specific calcium imaging, activity modulation, and brain-wide circuit mapping, we identify CeLC^PKCδ^ neurons as critical drivers of opioid withdrawal. Collectively, our data indicate that CeLC^PKCδ^ neurons are hyperactive and hypersensitive to noxious stimuli during opioid withdrawal, that reducing their activity can alleviate withdrawal, and that several input pathways to CeLC^PKCδ^ neurons may be directly modulated by opioid drugs via MORs.

Previous studies reported CeA cellular activity during opioid withdrawal ^25–27,29^. Our immunohistochemistry results (**Fig. 1**) and our photometry results (**Fig. 3**) extend these findings, evidencing rapid and sustained increases in the activity of CeLC^PKCδ^ neurons during fentanyl withdrawal. The most straightforward explanation for chronic fentanyl-induced changes in neural activity is that MORs and/or other opioid receptors are present in the circuitry that controls the activity of these neurons. Though MORs are Gi/o-coupled receptors, chronic MOR stimulation (*e*.*g*., by chronic fentanyl exposure) increases adenylyl cyclase expression and cAMP-mediated signal transduction, including depolarizing mechanisms, in MOR-expressing neurons ^78–82^. This paradoxical increase in signal transduction helps maintain basal signaling levels despite chronic MOR-mediated suppression of these processes. That compensatory increase in transduction and cellular activity can be “unmasked” when MOR stimulation either ends (*e*.*g*., spontaneous withdrawal) or is abruptly blocked (*e*.*g*., precipitated withdrawal) ^63,81,82^. This molecular process is called cellular dependence and represents a physiological state underlying physical dependence.

Adenylyl cyclase-dependent hyperactivity of MOR-expressing neurons, including CeLC^PKCδ^ neurons themselves or excitatory inputs to CeLC^PKCδ^ neurons, can explain both the increase in FOS and the sharp rise in Ca^2+^ activity that we detected during fentanyl withdrawal in CeLC^PKCδ^ neurons. Cooper et al. ^13^ found that approximately 40% of MOR lineage neurons express PKCδ protein; inversely, Chaudun et al^29^ found that ∼80% of *Oprm1*^+^ cells in the CeA also express *Prkcd* mRNA. Thus, there is a strong possibility that a subset of CeLC^PKCδ^ neurons can be directly modulated by chronic fentanyl via expression of fentanyl’s primary molecular target.

An alternative explanation for the heightened activity seen here is that MOR-expressing, and therefore opioid-sensitive, excitatory projections to the CeA are susceptible to cellular dependence. Their activity, in turn, could drive the activity of downstream CeLC^PKCδ^ neurons. Our parallel viral labeling strategies revealed that most brain areas with direct inputs to CeLC^PKCδ^ neurons contain MOR-expressing CeA projections (**Fig. 5**). The projections of several of these inputs are also involved in drug dependence and withdrawal, including those from the insular cortex ^69–71^, dorsal striatum ^72^, basolateral amygdala ^73,74^, and paraventricular thalamus ^75,76^. Cellular dependence within these input neurons provides a potential presynaptic mechanism for the hyperexcitability we see in the CeA during withdrawal.

While many of the areas we identified in our tracing studies are well-established loci of the effects of opioid drugs, the role of sensory cortical areas in opioid dependence may be underexplored. We found that both acute fentanyl administration and fentanyl withdrawal modulate the response of CeLC^PKCδ^ neurons to high-salience somatosensory stimuli (noxious hot water, airpuff; **Fig. 3**). Airpuff also has an auditory component, as the release of compressed air is accompanied by a loud puffing sound. These results suggest that opioids might affect the inputs from the areas of the brain responsible for processing these two sensations. Several areas our tracing approach identified participate in somatosensation and audition, including the somatosensory cortex, the secondary auditory cortex, and the temporal association area, as well as ventral posterior thalamic nucleus, part of the thalamic auditory system (**Fig. 5, Fig. S7**). In further support of sensory areas as sites of action of opioid drugs, emerging evidence shows a susceptibility of the mouse somatosensory and auditory cortex to persistent plasticity after perinatal opioid exposure ^83,84^. It is unknown whether such findings translate to animals exposed as adults, or whether *Oprm1*^+^ projections to the central amygdala are likewise affected. Alongside our results, the susceptibility of these sensory areas to perinatal opioid exposure warrants further investigations of *Oprm1*^*+*^ CeA projections from such regions in the context of opioid dependence.

Intriguingly, administration of a MOR antagonist animals with a prior injury induces a withdrawal-like cellular and behavioral phenotype ^13,63,82^. Doing so also reinstates hyperalgesia at the injury site, even well after the initial hypersensitivity has resolved – a phenomenon termed *latent pain sensitization* ^63,82^. Latent pain sensitization is also thought to result from cellular dependence, when injuries trigger constitutive activity of MORs that suppresses cellular processes producing hyperalgesia and hyper-sensitivity ^13,63,82^. The CeA specifically undergoes hyperactivity in this context, and the resulting pattern of latent sensitivity-related FOS occurs in a similar location in the CeLC as the withdrawalFOS we identified ^13^. Additionally, a large portion of latent sensitivity-induced FOS^+^ neurons co-express PKCδ. Thus, increased activity of CeLC^PKCδ^ neurons represents an overlapping mechanism of latent pain sensitization and opioid withdrawal. Overlap in the mechanism underlying both phenomena may explain the hyper-sensitivity patients often report during opioid withdrawal ^2,6,67,85^

We note that molecularly-defined neural populations in the central amygdala are not always homogeneous. Investigations of chronic pain-induced phosphorylated ERK and FOS find that they increase primarily in the posterior half of the CeA ^14^. In contrast, we found that the peak of *withdrawal*FOS occurs in the anterior half of the CeA, and that *withdrawal*FOS is elevated throughout the CeLC (**Fig. 1e**). Additionally, previous studies suggest that the CeA is a lateralized structure. Under this framework, the right CeA putatively dominates the processing and expression of aversion, and especially nociception ^12,16,24,48^, with the left CeA potentially suppressing nociception ^12^. We did not detect a robust effect of hemisphere in our studies, despite withdrawal causing a highly aversive state (**Fig. S3**). That our data diverge from these previous findings, but align with latent pain sensitization, indicates that there are multiple CeLC^PKCδ^ functional ensembles responding to aversive stimuli that may undergo plasticity under different conditions ^11,13,14^. Indeed, Kim et al ^37^ suggested that functional separation of PKCδ^+^ neurons exists along the anatomical axes of the CeA. Specifically, they found that CeC^PKCδ^ neurons in the anterior CeA receive different intra-amygdalar inputs, respond differently to aversive stimuli, and differentially drive behavior compared to CeL^PKCδ^ neurons located more posterior. Spatial transcriptomic analyses also show genetic divisions of CeLC^PKCδ^ neurons along the anatomical axes of the CeA ^30,31^. Here, we determined that *withdrawal*FOS peaks in the anterior-to-middle portion of the CeA, but is present throughout the anterior-posterior axis of the CeA (**Fig. 1**), suggesting overlap with both populations identified by Kim et al. Our viral injections targeted both CeC^PKCδ^ and CeL^PKCδ^ neurons in the anterior half of the CeA, centered around the peak of *withdrawal*FOS (**Fig. 1e, Fig. S3a, Fig. S6a**), so may have captured some of each “distinct” population. Thus, our results might mask some of their divergent roles. Future work could investigate this possibility using single-cell calcium imaging approaches or activity-dependent genetic capture (e.g., *Fos*TRAP) ^86^.

The aversive experience of opioid withdrawal makes abstinence challenging and increases the risk of relapse. The high comorbidity between persistent pain and Opioid Use Disorder suggests shared biological mechanisms, and our findings, which align with a model of latent pain sensitization, indicate that CeLC^PKCδ^ hyperactivity may be one such mechanism. By identifying the role of the nociception-activated and -responsive CeLC^PKCδ^ neurons in opioid withdrawal, our study reveals how a neural circuit involved in pain processing is affected by chronic opioid use. In total, these results underscore the need for cell-type-specific investigations within the CeA, which will be essential for developing targeted treatments for opioid use disorder that do not compromise pain management.

## MATERIALS AND METHODS

### Animals

All experiments described here were approved by the University of Pennsylvania Institutional Animal Care and Use Committee and performed in accordance with the National Institute of Health (NIH) guidelines for animal research. Male and female mice between 3-5 months of age were housed in a temperature- and humidity-controlled vivarium on a 12hr:12hr reverse light/dark cycle (lights off at 9:30AM). All experimental procedures took place during mice’s dark phase under red light. Mice were housed in groups of 2-5 prior the beginning of fentanyl treatment, at which point they were individually housed (for experiments in **Fig. 4** where dosage is reported) or housed in groups of 2-3 (all other experiments). Except where noted, all experiments were performed in transgenic *Prkcd*-Cre mice (Tg(Prkcd-glc-1/CFP,-cre)EH124Gsat, MGI ID: 3844446), which were obtained from Drs. Matthew Hayes and Tito Borner. For experiments in **Figs. 5d-f + Fig. S9**, *Oprm1*-Cre mice were donated by Dr. Richard Palmiter (B6.Cg-Oprm1tm1.1(cre/GFP)Rpa/J, Jackson Laboratories, strain #035574). All mice in the present studies were bred inhouse from a male mouse heterozygous for the Cre transgene crossed to a female C57bL/6J mouse (Jackson Laboratories, strain # 000664). Experiments included heterozygous male and female offspring from these crosses. Food and water were available *ad libitum* throughout all experiments; during fentanyl treatment, food and fentanyl-treated water was available *ad libitum*. All data collection involving mouse handling was performed by female experimenters (LMW, JWKW, AYJ, or AML) ^45^.

### Experimental model of fentanyl dependence

The standard water bottle for each cage was replaced with a 15 mL Amuza Drink-O-Measurer bottle with a double ball-bearing sipper tube, marked by the manufacturer in 1 mL increments (Amuza Drink-O-Measurer). On day 0, the water of fentanyl-drinking mice was replaced with animal facility tap water treated with 0.02 mg/mL fentanyl citrate (Covetrus, item #055012), modified after previous studies using fentanyl-treated water as a means of unsignaled drug delivery (**Fig. 4a**) ^46,47^. Water-drinking control mice drank untreated facility tap water from identical bottles. For experiments where intake is reported, mice were weighed daily at 1:00 PM, and the intake volume was measured to the nearest 0.5 mL to calculate daily consumed dose. Bottle solutions were replenished daily. For experiments where intake is not reported, bottle volume was checked daily and replenished as needed. After 8 days, all bottles were replaced with standard animal facility water bottles containing untreated water either immediately following opioid receptor antagonist administration (**Figs. 1 and 3**) or 20-24 hours prior to spontaneous withdrawal assessments (**Fig. 4**).

### Drug Preparation

For oral fentanyl administration, fentanyl citrate (50 ug/mL, Covetrus, item #055012) was diluted in animal facility tap water to a final concentration of 0.02 mg/mL and stored at room temperature for no more than 4 weeks. Fentanyl citrate was diluted in 0.9% sterile saline and delivered at a dose of 0.2 mg/kg for acute treatment in **Fig. 3**. Naltrexone hydrochloride (Hellobio, cat # HB2452) and naloxone hydrochloride (Hellobio, cat #HB2451) were dissolved in saline and stored at 4°C for up to a year. Meloxicam (Loxicom, Midwest Veterinary Supply, cat #515.50000.3) was diluted in saline and stored at room temperature for up to a month. Pentobarbital (Fatal Plus containing 390 mg/mL pentobarbital, Covetrus, cat # 035946) was stored at room temperature for up to a year and delivered undiluted at a volume of 0.08 mL. Unless otherwise noted, all drugs were delivered via subcutaneous injection at a volume of 10 mL/kg.

### Viral vectors

Viral vectors were diluted to their final working concentration in sterile phosphate-buffered saline (PBS), where necessary. Vectors were stored in 10 uL aliquots of their working titer at −80°C prior to use; during use, aliquots were stored at 4°C for up to 5 days. All viruses were injected at a flow rate of 100-125nL/min to the right central amygdala (stereotaxic coordinates: anterior-posterior AP −1.05 or −1.10 mm, medial-lateral ML +2.78 mm, dorsal-ventral DV −4.82 mm relative to bregma) or the right parabrachial nucleus (stereotaxic coordinates: AP −5.33mm, ML +1.10mm, DV −3.65mm relative to bregma) due to previous reports of lateralized function of the central amygdala in response to aversive stimuli ^12,16,48^. For photometry experiments, 500 nL of AAV9-hsyn-FLEX-GCaMP6f (titer: 2.8×10^12^ vg/mL; Addgene, Plasmid #100837) was injected into the right CeA of *Prkcd*-Cre mice. For chronic inhibition experiments, 300 nL of AAVDJ-CMV-Kir2.1-zsGreen (titer: 1.35×10^12^ vg/mL; from Dr. Marc Fuccillo) or AAV5-hsyn-DIO-EGFP (titer:1.3×10^12^ vg/mL; Addgene, Plasmid #50457) was injected in *Prkcd*-Cre mice. For monosynaptic input tracing experiments, AAV2-hsyn-TCB-mCherry (1.7×10^12^ cfu/mL; from Dr. Kevin Beier) and AAV8-hsyn-G (1.0×10^12^ cfu/mL; from Dr. Kevin Beier) were mixed 1:1 and injected into the right CeA of *Prkcd*-Cre mice at a volume of 500 nL. RABV*d*G-GFP (8×10^8^ cfu/mL; from Dr. Kevin Beier) was injected 3 weeks later at the same location, at a volume of 500 nL. For labeling of *Oprm1*^+^ and *Oprm1*-inputs to the CeA, AAVrg-EF1a-Nuc-fl(mCherry)-EGFP was injected at a volume of 250 nL in *Oprm1*-Cre mice (2.5×10^13^ vg/mL; Addgene, Plasmid #112677).

### Two-bottle choice

Mice were individually housed in standard cages with a 3D-printed two-bottle choice apparatus equipped with two 15 mL conical tubes modified as sipper bottles ^49^. One tube contained untreated tap water from the animal facility, and the other contained the same water treated with 0.02 mg/mL fentanyl citrate. The weights of the mice and the intake volume from each bottle were recorded daily at 12:00 pm for 16 days. The positions of the bottles were switched every day to minimize side preference effects.

### Immunohistochemistry

Formalin-fixed tissue was coronally sectioned at 30-50 um, protected from light, and stored at 4°C in PBS until free-floating IHC was performed. Tissue was then washed in 0.5% Triton-X100 detergent in PBS (Sigma-Aldrich) 4 times for 10 min each to permeabilize the cell membranes, rinsed 4 times for 10 min in fresh PBS, and incubated in normal donkey serum blocking solution (Jackson ImmunoResearch, 10% in 0.1% Triton-X+PBS) to block off-target antibody binding. Slices were then incubated in the primary antibody diluted in blocking solution to the desired concentration for 48 hrs at 4°C. Primary antibodies include: rabbit anti-FOS (1:1000, Synaptic systems, cat # 226008), guinea pig anti-FOS (1:1000, Synaptic systems, cat # 226308), mouse anti-PKCδ (1:1000, BD Transduction Laboratories, cat # 610397), rabbit anti-MOR (1:1000, Abcam, cat # ab134054); chicken anti-GFP (1:1000-1:2000, Abcam, cat # ab13970), rabbit anti-dsRed (1:1000, Takara, cat # 632496). Slices were then washed in PBS 4 times for 10 min, followed by incubation with secondary antibody diluted in blocking solution to the desired concentration for 24 hrs at 4°C. Secondary antibodies include: Alexafluor Donkey anti-mouse 647 (1:500, Invitrogen Thermo Fisher, cat # A31571), Alexafluor Donkey anti-guinea pig 647 (1:500, Jackson ImmunoResearch, cat # 705-605-148), Alexafluor Donkey anti-rabbit 488 (1:500, Invitrogen Thermo Fisher, cat # A21202), Alexafluor Donkey anti-mouse 647 (1:500, Invitrogen Thermo Fisher, cat # A31571), Alexafluor Donkey anti-chicken 488 (1:500, Jackson Immunoresearch, cat # 703-545-155), Alexafluor Donkey antichicken 594 (1:500, Jackson ImmunoResearch, cat # 703-585-155). Tissue was then washed in PBS 4 times for 10 min, then mounted onto glass slides and incubated with a 1:10,000 solution of DAPI counterstain diluted in water for 10 min (Fisher Scientific). The tissue was then covered with Fluoromount-G mounting medium (Fisher Scientific), coverslipped, and dried flat overnight in a dark, cool place before imaging.

### Fluorescent *in situ* hybridization

The RNAScope Multiplex Fluorescent v2 Assay (Advanced Cell Diagnostics) was used to label mRNA according to the manufacturer’s instructions. Briefly, mice were perfused as described previously, and brains were removed and postfixed in 10% normal buffered formalin for 24 hrs. The whole brains were then submerged in 10%, 20%, then 30% sucrose at 4°C until the tissue sank to the bottom of the container (approximately 24-48 hrs for each concentration). Brains were then frozen in TissueTek O.C.T. compound (Thermo Fisher Scientific) at −20°C. 16 um sections of the CeA or PBN were collected and placed onto dry SUPERFROST PLUS (Thermo Fisher Scientific) slides while still frozen. Slides were either processed immediately or stored at −80°C for up to 3 months. Slides were next washed with PBS for 1 min, then baked for 30 min at 60°C in a HybEZ oven (Advanced Cell Diagnostics). Slides were postfixed in chilled 10% neutral buffered formalin for 15min at 4°C. The slides were then dehydrated by immersion in 50%, then 70%, then 100% EtOH x 2 for 5 min at room temperature. The slides were incubated in 6% hydrogen peroxide for 10 min at RT, then washed several times in distilled water, then pretreated with RNAScope Target Retrieval Reagent heated to 99°C for 15 min, inside a commercial steamer (Oster). Slides were rinsed for 15 sec, transferred to 100% ethanol for 3 min, and dried. A hydrophobic barrier was drawn around the tissue using an Immedge pen, and the slides were dried overnight in a closed drawer at room temperature. The following day, slides were incubated in the provided Protease III reagent for 30 min at 40°C and washed twice with distilled water. Slices were next incubated in the desired mixture of cDNA hybridization probes for 2 hrs at 40°C, then washed twice in the provided Wash Buffer. Probes used included: Mm-Oprm1-C1 (Advanced Cell Diagnostics, Cat # 315841), Mm-EGFP-C2 (Advanced Cell Diagnostics, Cat #400281-c2), 4-Plex Negative control (Advanced Cell Diagnostics, Cat#321831). Amplification was performed by incubating tissue at 40°C in the provided FL v2 Amp1 (30 min), FL v2 Amp2 (30 min), then the FL v2 AMP3 (15 min), with 2 min washes in the provided wash buffer before and after each step. Slices were then fluorescently stained by first incubating sections in the provided TSA Plus HRP solution for the desired channel for 15 min at 40°C, then in a 1:5000 solution of Opal 520 or 690 fluorescent dye (Akoya Biosciences, cat # FP1487001KT; cat # FP1497001KT) diluted in the provided TSA buffer for 30 min at 40°C, then a final incubation with the provided HRP blocker for 15 min at 40°C, with 2 min washes in the provided Wash Buffer between each incubation. This process was repeated for each channel. Lastly, sections were counterstained using the provided DAPI reagent, mounted with Fluoromount-G mounting medium, and coverslipped. Pairs of sections were collected such that for each stained section, an additional section was hybridized with the 4-Plex Negative control probe, which contained bacterial RNA instead of a cDNA.

### Imaging and quantification

All imaging was performed using a Keyence BZ-X800 all-in-one fluorescent microscope with PlanApo-λ x4, PlanApo-λ x20, and PlanApo-λ x40 objectives. Image processing was completed for all images using the Keyence BZ-X analyzer software (version 1.4.0.1). For placement checks/viral transduction and targeting verification, slices were imaged in a single plane of focus at 4x magnification. For histological quantification, slices were imaged with 20x-magnified z-stacks or, for fluorescence *in situ* hybridization, in a single plane of focus at 40x magnification. The exposures for FITC, TRITC, and Cy5 were adjusted to avoid overexposed pixels. Exposures were standardized such that a single exposure for each channel was used across all tissue in the same experiment. For quantification of cells with a single marker protein or non-overlapping markers, quantification was performed manually using the Adobe Photoshop Counter function or semi-manually using Cellpose 2, as indicated^50,51^. When quantifying colocalization of 2 or more markers, quantification was performed semi-manually using HALO-AI software (Indica Labs). When using Cellpose 2 or HALO, a single set of settings was developed and used to quantify all tissue in the same experiment.

### Stereotaxic surgery

Adult mice (∼12 weeks of age) were anesthetized with isoflurane gas in oxygen (induction: 5%; maintenance: 1.0-2.5%) and mounted onto a stereotaxic frame (KOPF instruments). Mice were administered meloxicam (5 mg/kg, s.c.) and 1 mL saline (s.c.) at the beginning of the surgery. Eye lubricating gel (Optixcare) was applied at the beginning of surgery and every 45 min throughout, and reflexes and respiratory rates were assessed every 10 minutes to ensure a surgical plane of anesthesia. The skull was manually leveled and a ∼1 mm craniotomy was drilled over the target area. A 10 uL Hamilton Nanofil syringe fitted with a 33 gauge beveled needle was slowly lowered into the right CeLC (coordinates from Bregma: AP −1.05mm or −1.10mm, ML +2.78mm, DV −4.82mm) or the right parabrachial nucleus (coordinates from Bregma: AP −5.33mm, ML +1.10mm, DV-3.65mm), and 250-500 nL of AAV or RABV*d*G vectors containing transgenes of interest were infused at a rate of 100-125nL/min. All injections were performed unilaterally in the right hemisphere because of previous reports of lateralization in central amygdala function in response to aversive stimuli and in the maintenance of aversive states ^16,24,48,52^. Incisions were closed with 5-0 Vicryl sutures (Ethicon).

For fiber photometry experiments, immediately after virus injections the skull was scored with a scalpel blade and 2 small screws (∼1.7 mm diameter, 1.5 mm length) were placed distal to the craniotomy as additional anchor points. Immediately after the viral injection, an optical fiber (5.0mm fiber, 400 um diameter, Doric Lenses) was slowly lowered to approximately 0.2mm over the target coordinate (in the DV plane) and fixed to the skull using MetaBond (Parkell) followed by Jet Set dental acrylic (Lang Dental) to create a reinforced headcap and cover exposed skull. The dental acrylic was mixed with black iron oxide power to reduce the chance of any excitation or emission light leak.

Following all surgeries, mice recovered under a heat lamp until they fully regained the righting reflex, and were then returned to their homecage. Mice were monitored daily for 3 days following surgery to ensure integrity of the sutures and headcaps. Meloxicam was administered again 24 hrs after surgery for viral injections and 24, 48, and 72 hrs after fiber implant to minimize pain and brain inflammation

### Behavioral testing

*Prkcd*-Cre mice were used in all behavior experiments, and both males and females were included in all experimental groups. Mice were habituated to behavioral apparatuses for 1 hr each for the 2 days prior to the beginning of behavioral testing, and habituated to the apparatus for 30 minutes with the experimenter present before beginning testing each day. Mice were returned to the colony room after all animals in the group had completed behavioral testing each day. Males and females were not tested in the same room at the same time. All experiments took place between 10:30AM-6:30PM.

#### Apparatus

For all experiments, mice were placed in individual enclosures on a custom-made elevated rack with a perforated metal floor (61 cm x 26 cm; **Fig. S4i**). For experiments conducted in **Figs. 1 and 4**, the enclosures were 16 cm x 16 cm x 38 cm height plexiglas rectangular enclosures with a black foam lid. For fiber photometry experiments (**Fig. 2, 3)** the enclosures were 7 cm diameter x 10 cm height red Plexiglas cylinders with a custom-printed, form-fitting lid containing a passage for the photometry patch cable. These apparatuses accommodated both withdrawal scoring and nociceptive testing without requiring movement of the animal. Unless otherwise indicated, all stimuli were applied to the left hind-paw, contralateral to the right CeLC viral injection.

#### Spontaneous withdrawal assessment

Precipitated withdrawal was assessed immediately following antagonist injection; spontaneous withdrawal was assessed 20-24 hrs following the removal of fentanyl-treated water **(Fig. 4a)**. A Logitech webcam (C920x HD Pro) protected by a custom Plexiglas enclosure (58 cm x 32 cm x 20 cm) were used to record somatic signs from the underside of the mouse (**Fig. S4i**); videos were scored offline using BORIS behavioral software^53^ by an experimenter blinded to experimental condition. To score each video, mice were continuously observed for 15min and the total instances (>1sec) of genital grooming, digging at the corners of the apparatus, bouts of running, jumping, paw tremors, head shakes, wet dog shakes, and the total number of observed fecal boli were recorded. In addition, the presence/absence of a resting tremor and teeth chattering were recorded once for each minute they were observed. All instances and presence of events (1 point per minute present) were summed to get the global withdrawal score for each animal. In Supplemental Fig. 4 and 5, behaviors were further divided into three subscores: an autonomic score (sum of fecal boli and minutes with teeth chattering), affective score (genital grooming, running, jumps, digging) and shakes/tremors score (paw tremors, head shakes, wet dog shakes, minutes with resting tremor). Ethovision XT (Noldus, version 15) was run on the same 15 min video to calculate the distance traveled for each mouse.

#### Von Frey Up-down

TouchTest sensory evaluator filaments (North Coast Medical and Rehabilitation Products) of logarithmically increasing thickness (range: 0.07-6 g, initial filament: 0.16 g) were sequentially applied to the left hindpaw until they bent slightly. The force necessary to produce a hindpaw withdrawal response was calculated using the Up-Down method as described previously^54,55^ and reported as 50% withdrawal threshold.

#### Warm-water tail withdrawal

A Sous-vide Precision Cooker Nano 2.0 (Anova) maintained a water bath at 45°C, 48°C, or 50°C; we chose these temperatures to engage different combinations of transient receptor potential channels that control innocuous and noxious heat sensation ^56^. Mice were removed from the apparatus and briefly restrained in a cotton towel. The distal 1-2 cm of their tail was submerged in the water bath. The tail was immediately removed once it curled or flicked, and the latency to do so was recorded. If the mouse did not withdraw its tail within 20 sec, it was removed to avoid tissue injury, and the latency was recorded as 20 sec. The mouse’s tail was dried with a paper towel, and the mouse was returned to the apparatus. Temperatures were tested from lowest to highest, and a minimum of 15 min elapsed between consecutive tests.

#### 55°C hot water drop

A Fisher Micro hotplate (Thermo Fisher Scientific) was used to maintain approximately 50 mL of water at 55°C. Water droplets were expressed from the end of a 1 mL syringe and applied to the plantar surface of the left hindpaw while the mouse was on the apparatus. The mouse’s response was video-recorded and scored offline using BORIS behavioral software^53^ by an experimenter blinded to experimental condition. Total time spent in affective behaviors (attending the paw, licking the paw, guarding the paw, rearing, running, and jumping) were scored for the 30 sec immediately following the hot water application

#### 50°C inescapable hotplate

Mice were removed from the apparatus and placed on an enclosed hotplate (Bioseb hot and cold plate; 16.5 cm x 16.5 cm floor) maintained at 50°C. The mice were promptly removed after 60 sec to avoid tissue damage and immediately returned to their home cage. Cameras (Logitech C920x HD Pro Webcam) were set up on either side of the hotplate to videorecord behavior while the mouse moved freely on the plate. Videos were scored offline using BORIS behavioral software^53^ by an experimenter blinded to experimental condition. Instances of paw flicks, hops, and jumps, and the duration of attending behaviors (licking, attending, or guarding of one or more paws), and escape behaviors (rapid rearing, running, hopping, and jumping), were scored for the full 60 sec hotplate exposure.

### *withdrawal* FOS induction and immunohistochemistry

Male and female *Prkcd*-Cre mice were individually housed on day 0 of fentanyl or water administration. On Treatment Days 6 and 7, mice were habituated in their home cages to the testing room for 2 hrs daily. On Day 8, mice were habituated in their home cages to the testing room for a minimum of 30 min. Mice were then injected with naltrexone (1 mg/kg, s.c.) or saline and immediately returned to their homecage. The fentanyl-treated water was removed and replaced with untreated water. We waited 15 min for naltrexone to take effect, then an additional 90 min for the peak of FOS protein production. Thus, 105 min after naltrexone or saline injection, mice received an overdose of pentobarbital (Fatal Plus, 0.08 mL, i.p.) and were placed back in their homecage until toe-pinch and corneal reflexes were absent. Mice were then promptly transcardially perfused with phosphate-buffered saline followed by 10% normal-buffered formalin. The brain was removed and postfixed in 10% normal-buffered formalin overnight, then incubated in 30% sucrose in PBS until the tissue sunk. 30 um coronal slices through the expanse of the central amygdala were collected on a cryostat (CM3050S, Leica Biosystems). Every 3^rd^ 30 um slice was stained for FOS and PKCδ or GFP using immunohistochemistry (see section for details). The anterior-posterior coordinate was determined for each slice and borders were drawn based on the Paxinos and Watson 5^th^ edition stereotaxic atlas ^57^, and the CeA was imaged at 20x magnification with z-stack. FOS^+^ and/or PKCδ+ neurons were quantified in each of the CeA subnuclei from each side of one slice per 0.2 mm. Quantification was performed using CellPose 2.0 (**Fig. 1a-e, Fig. 5e, Fig. S1**)^50,51^ for single-fluorophore images, or using HALO-AI software (Indica Labs; **Figs. 1f-i, Fig. S9f-g**) when quantifying colocalized markers. Except where noted, each point represents the sum of the cells in the left and right hemisphere (*78, 79*).

### In vivo Fiber photometry

500 nL of AAV9-hsyn-FLEX-GCaMP6f-WPRE-SV40 was injected into the right CeA of male and female *Prkcd-*Cre mice (AP −1.10, ML +2.78, DV −4.82) and a 400 um borosilicate fiber-optic cannula was implanted 200 um above the injection site (5 mm length, 0.66 NA; Doric Lenses; see surgery section for details). A minimum of 3 wks of surgical recovery elapsed between surgery and the beginning of habituation. For two days prior to the first test day, and on each test day, mice were habituated to the testing room in their home cage for 1 hr without the experimenter present, then to the apparatus and patch cord for 15 min with the experimenter present.

Optical recordings of GCaMP6f fluorescence were acquired using an RZ10x fiber photometry detection system (Tucker-Davis Technologies), a processor with Synapse software (Tucker-Davis Technologies), and optical components (Doric Lenses and ThorLabs). LED-generated light was filtered through a fluorescence minicube at spectral bandwidths of 460 nm and 405 nm and passed through a pre-bleached, low auto-fluorescence mono fiber-optic patch cord (Doric Lenses) connected to the external portion of each mouse’s fiber-optic cannula via a zirconia mating sleeve (Doric Lenses). The power output at the tip of the patch cable was adjusted daily to ∼50uW for the 460 nm channel (calcium-dependent signal; modulated at 210 Hz), and to ∼15uW for the 405 nm channel (isosbestic control; modulated at 330 Hz). Signals were low-pass filtered at 6 Hz. All recordings utilized both channels.

On the first (baseline) test day, 4 stimuli were presented to the mouse for 10 trials each with a 90 sec intertrial interval (ITI): 0.16 g von Frey Filament (TouchTest), 25-gauge needle “pinprick”, 55°C hot water drop, all delivered to the left hindpaw; and an airpuff from a condensed air canister delivered 1 cm from the left cheek/eye (**Fig. 2a**). A TTL pulse was delivered to the Synapse software by the experimenter at the moment of stimulus application by means of a custom-built trigger button. Trials were aborted if the mouse responded behaviorally to the approach of the stimulus. Additionally, trials were excluded if the stimulus did not contact the mouse or produce the expected behavioral responses. Between each stimulus type, the recording was turned off for 5 min to minimize photobleaching.

On both the acute fentanyl and withdrawal test days, fluorescence was recorded for 7 min to establish a baseline period prior to injection (**Fig. 3a**). While continuously recording, the mouse was removed and injected with either fentanyl (0.2 mg/kg, s.c.) or 0.9% saline for the acute fentanyl test in naïve mice (Fig. 4B), or naloxone for the withdrawal test (3 mg/kg, s.c), and returned to the enclosure, all while still attached to the patch cord. A TTL pulse was delivered immediately before the removal of the mouse, and immediately after the return of the mouse to the enclosure (the “injection period”). Fluorescence was then recorded for 15 min. Following a 5 min rest in the recording, mice were again presented with the highly-salient stimuli, the 55°C hot water droplet and the airpuff. To ensure all testing occurred while fentanyl/naloxone was at its peak effectiveness, only 5 trials of each stimulus were delivered, again with a 90 sec ITI. At the end of the last recording, mice were immediately disconnected from the patch cable and returned to their homecage.

At the completion of experiments, all mice were perfused and immunohistochemistry was performed to amplify GCaMP6f fluorescence using chicken anti-GFP and donkey anti-chicken AF488 antibodies. Each brain was imaged, and animals without virus expression and proper fiber placement in the CeA were excluded from the experiment. In addition, two animals were excluded only from the withdrawal recording: one mouse in the fentanyl-drinking group was excluded and euthanized because of a sudden health decline; and one animal in the water-drinking group was excluded from the withdrawal recording *only* because after data processing it was apparent that the patch cord had lost contact the fiber during the naloxone injection. This was fixed prior to the stimulus tests.

### Fiber photometry data

Data from the Synapse software was processed using an independent deployment of the open-source pMAT fiber photometry analysis software and MATLAB^58^. For stimulus-locked recordings, peri-event time histograms were generated in pMAT from −10 sec to 10 sec relative to the TTL pulse for each trial, with a baseline sampling window from −10 sec to −5 sec, and both the trial-based dF/F and z-scores were exported. All trials of a given stimulus were averaged for a given mouse; therefore, each data point is one mouse. The max amplitude of the Ca^2+^ responses in the first 5 sec following the stimulus (peak z-score) were determined from the mean trace, and the net AUC of the mean dF/F trace or z-scored dF/F was calculated in Graphpad Prism for the 5 sec following stimulus presentation. These AUCs were statistically compared to baseline areas under the curve calculated from −10 sec to −5 sec relative to the stimulus, to minimize the influence of Ca^2+^ responses related to the experimenter approaching the mouse. For the withdrawal tests, we further adjusted the ∼25 min drug injection recording to account for the persistent downward trend of the signal due to photobleaching. The entire recording and the timestamps for the TTLs around the injection were exported into MATLAB, the signals of each channel was smoothed and downsampled from 10 Hz to 3 Hz, and the 465 nm signal recording for the pre-drug baseline was re-fit to the same period of the smoothed 405 nm isosbestic recording using custom MATLAB code. The dF/F was re-calculated from this smoothed and downsampled signal, and a baseline z-score for each timepoint was calculated from this data using the 7 min pre-injection period as the baseline window.

### Viral-mediated overexpression of Kir2.1

300 nL of AAVDJ-CMV-DIO-Kir2.1-zsgreen or AAV5-hsyn-DIO-EGFP was injected into the right CeA of male and female *Prkcd*-Cre mice. 2.5-3 wks later, mice underwent baseline testing to determine values for mechanical response thresholds (von Frey updown), thermal reflexive thresholds (warm water tail withdrawal at 45°C, 48°C and 50°C), and responses to a 55°C hot water drop applied to the hindpaw, all as described above (**Fig. 4g**). After a 30min period to habituate to the testing apparatus, mice underwent the von Frey up-down procedure on the left, then right paw. Next, a hot water drop was applied to the left, then right paw. Tail flick latencies were then collected in order of increasing temperature. A minimum of 15 min elapsed between each trial. Mice were returned to their homecage immediately after the 50°C tail withdrawal test. After baseline testing, mice began the 8-day fentanyl or water drinking procedure. All Drink-o-measurer bottles were removed late afternoon on the 8^th^ drinking day and replaced with regular animal facility water bottles. On Day 9, 20-24 hrs after bottle removal, mice were returned to the same testing room and were tested on all nociceptive procedures in the same sequence as on the baseline day. Video recordings of the mice were also collected during the habituation period, and behavior from the first 15 min of the period were scored offline to obtain the global withdrawal score and distance traveled. Following the conclusion of the experiments, mice were perfused, and 50 um slices were collected through the anterior-posterior axis of the CeA of each animal to confirm viral transduction and targeting. Data from mice in which few cells were visible in the CeA or with significant additional off-target expression were excluded.

For experiments assessing the ability of Kir2.1 in CeLC^PKCδ^ neurons to reduce withdrawalFOS, we injected a separate group of *Prkcd-*Cre mice with AAVDJ-CMV-DIO-Kir2.1-zsGreen or AAV5-hsyn-DIO-EGFP. 3 weeks later, mice were individually housed and the water bottles of all mice were treated with 0.02 mg/mL fentanyl. On the eighth day of fentanyl treatment, mice were moved in their home cage to a procedure room and habituated for 1hr with the experimenter present. Mice were then removed from their home cages and promptly injected with 1 mg/mL naltrexone (s.c.), returned to their homecage, and their water bottles were replaced with untreated water. 105 minutes later, mice were injected with pentobarbital (Fatal Plus, 0.08 mL, i.p.) and immediately removed from the procedure room. Once reflexes were absent, the mouse was perfused and tissue was collected and stained as described previously using rabbit anti-FOS and donkey anti-rabbit 647 antibodies (see FOS induction and immunohistochemistry sections for more detail). Because only minimal GFP was visible in unstained tissue, and because we did not plan to directly compare the total number of green fluorescent cells between the Kir2.1-injected mice and the GFP-injected mice, 30 um slices of the GFP-injected mice were additionally stained with chicken anti-GFP and donkey anti-Chicken 488 antibodies.

### Monosynaptic rabies tracing

500 nL of a 1:1 mixture of AAV2-DIO-TCB-mCherry and AAV8-FLEX-RABV*d*G was injected into the right CeA of male and female *Prkcd*-Cre mice. Two weeks later, the same area was injected with 500 nL of RABV*d*G-GFP. Five days later, mice were perfused and the brains were postfixed and cryoprotected. Consecutive 50 um slices were collected from the entire brain. Every 2^nd^ 50 um slice from the central amygdala was imaged at 20x to confirm the presence of putative starter cells. We then selected areas of interest from an initial scan of the tissue. We imaged the right hemisphere of every 2^nd^ slice from +0.70 mm AP to −4.20 mm AP, and from −4.70 mm to −5.80 mm AP at 20x magnification with z-stacks. GFP+ input cells were counted manually in FIJI ^59^ using the Cell Counter tool, and the number of cells expressing both mCherry and GFP (i.e. putative starter cells) were counted in every 2^nd^ CeA slice using HALO-AI. The x-y coordinates of each identified GFP+ cell were exported from FIJI, and cells/slices were aligned to the Kim Lab Unified Anatomical Atlas^60^ using the open-source, MATLAB-based atlas registration program SHARCQ ^61^. Cells that were identified in individual subnuclei of brain structures were summed to result in a single number for each macrostructure according to the headings present in the Unified Anatomical Atlas.

In a separate group of RABV*d*G-injected mice, 50 um slices were collected from the central amygdala to verify starter cell expression, then 5 pairs of 16 um slices were collected from the parabrachial nucleus of each mouse. RNAScope was performed as described, to stain for *Oprm1* mRNA (using Mm-Oprm1-C1 cDNA probe and Opal 690 fluorescent dye) and to simultaneously amplify the EGFP in the RABV*d*G vector (using Mm-EGFP-C2 cDNA probe and Opal 520 fluorescent dye). The number of GFP+ input cells with and without *Oprm1* mRNA colocalization in the right PBN were manually quantified using the Counter tool in Photoshop (Adobe).

### Identification of opioidergic inputs to the central amgydala

250 nL of AAVrg-EF1a-Nuc-fl(mCherry)-EGFP was injected into the right CeA of male and female *Oprm1-*Cre mice. Eight weeks later, mice were administered pentobarbital (Fatal Plus, 0.08 mL, i.p.) and monitored until the toe pinch and corneal reflex were absent. Mice were then transcardially perfused, brains were dissected and post-fixed, and 50 um slices were collected every 2 mm throughout the brain. Both the mCherry and GFP fluorescent signals were amplified using immunohistochemistry (antibodies: rabbit anti-dsRed and donkey anti-rabbit 594; chicken anti-GFP and donkey anti-chicken 488). Regions we had previously identified as having dense inputs to CeLC^PKCδ^ neurons with the RABV*d*G study were imaged as 20x-magnified z-stacks.

In a separate group of *Oprm1*-Cre mice, 500 nL of AAV5-hsyn-DIO-EGFP was injected into the right PBN (−5.4mm AP, +1.35mm ML, −3.65mm DV). Mice in this study also received a unilateral injection of AAV9-hsyn-DIO-mCherry in the right CeA (not shown). 6 weeks later, mice were administered pentobarbital (Fatal Plus, 0.08 mL, i.p.) and monitored until the toe pinch and corneal reflex were absent. Mice were then transcardially perfused, brain tissue was collected and post-fixed, and every 2^nd^ 50 um slice was collected through the CeA and the PBN. Representative images were taken of putative PBN^MOR^ cell bodies in the PBN (i.e., EGFP+ neurons) and EGFP+ terminals in the CeA as 20x-magnified z-stacks.

### Statistical analyses

Power analyses were conducted in G*Power^62^ at the beginning of each experiment to determine minimum sample sizes and adjusted as needed based on observed effect sizes. Male and female mice were used in every experiment. Unless otherwise noted, data are presented as the mean +/-standard error of the mean, with gray circles representing the observed value for an individual subject. Kolgomorov-Smirnoff tests were used to assess the normality of data sets, and two-tailed paired or unpaired t-tests, Mann-Whitney tests, One- and Two-Way ANOVAS, or mixed model analyses were calculated as indicated using Prism software (Graphpad, v9). Paired t-tests or repeated-measures ANOVAs were used when comparing subjects’ results to their own baselines. Where ANOVA or mixed model analyses showed significant main effects or interactions, p-values for the relevant comparisons were adjusted using Bonferroni’s method.

## ACKNOWLEDGMENTS

We thank Dr. Nicholas Betley (Penn), Dr. Julie Blendy (Penn), Dr.Yarimar Carrasquillo (NICCIH), and Dr. Heath Schmidt (Penn) for input on experimental design. We thank Corder Lab members past and present for invaluable feedback on the project, especially to Dr. Blake Kimmey, Dr. Nora McCall, and Leann Goldberg for conceptual and technical assistance. We thank Dr. Matthew Hayes (Penn), Dr. Tito Borner (Penn), Dr. Richard Palmiter (University of Washington), Dr. Kevin Beier (University of California, Irvine), and Dr. Marc Fuccillo (Penn) for contributing key resources. Finally, we thank the University Laboratory Animal Resources and veterinary staff at Penn for caring for and maintaining the Corder Lab mouse colony.

## Funding

National Institute of General Medical Sciences, DP2GM140923 (GC)

National Institute on Drug Abuse, R01DA056599 (GC)

National Institute on Drug Abuse, R01DA054374 (GC)

Alkermes Pathway Grant (GC)

Brain and Behavior Research Foundation NARSAD Young Investigator’s Award (GC)

National Institute on Drug Abuse, F31DA057795 (LMW)

National Institute on Drug Abuse, T32DA028874 (AYJ)

National Institute of Mental Health, R25MH119043 (AML)

## Author contributions

Conceptualization: LMW, GC

Methodology: LMW, AJ, GC

Formal analysis: LMW, AJ, MM

Investigation: LMW, AJ, JWKW, MZ, AML, MM, SAC

Writing - Original Draft: LMW, GC

Writing - Review & Editing: LMW, AJ, JWKW, GC

Visualization: LMW, GC; Funding: LMW, GC

Supervision: LMW, GC

## Competing interests

The authors declare that they have no competing interests.

## Data and materials availability

All data are available in the main text or the supplementary material

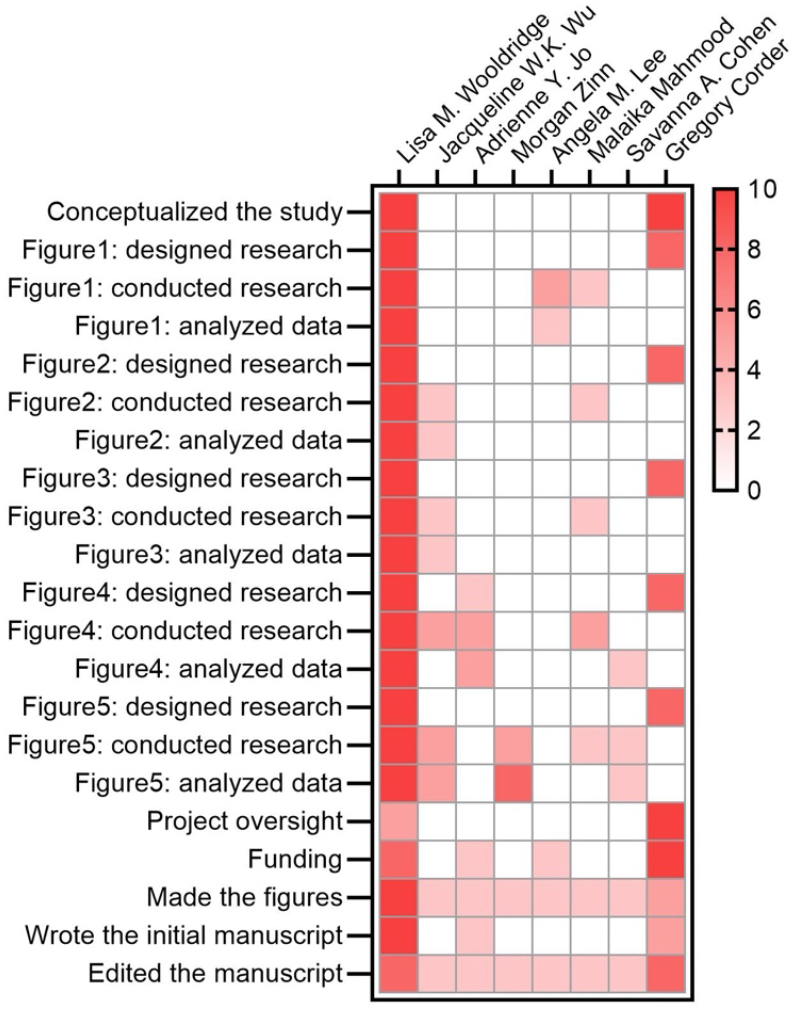

---

**Supplementary Fig. 1.**
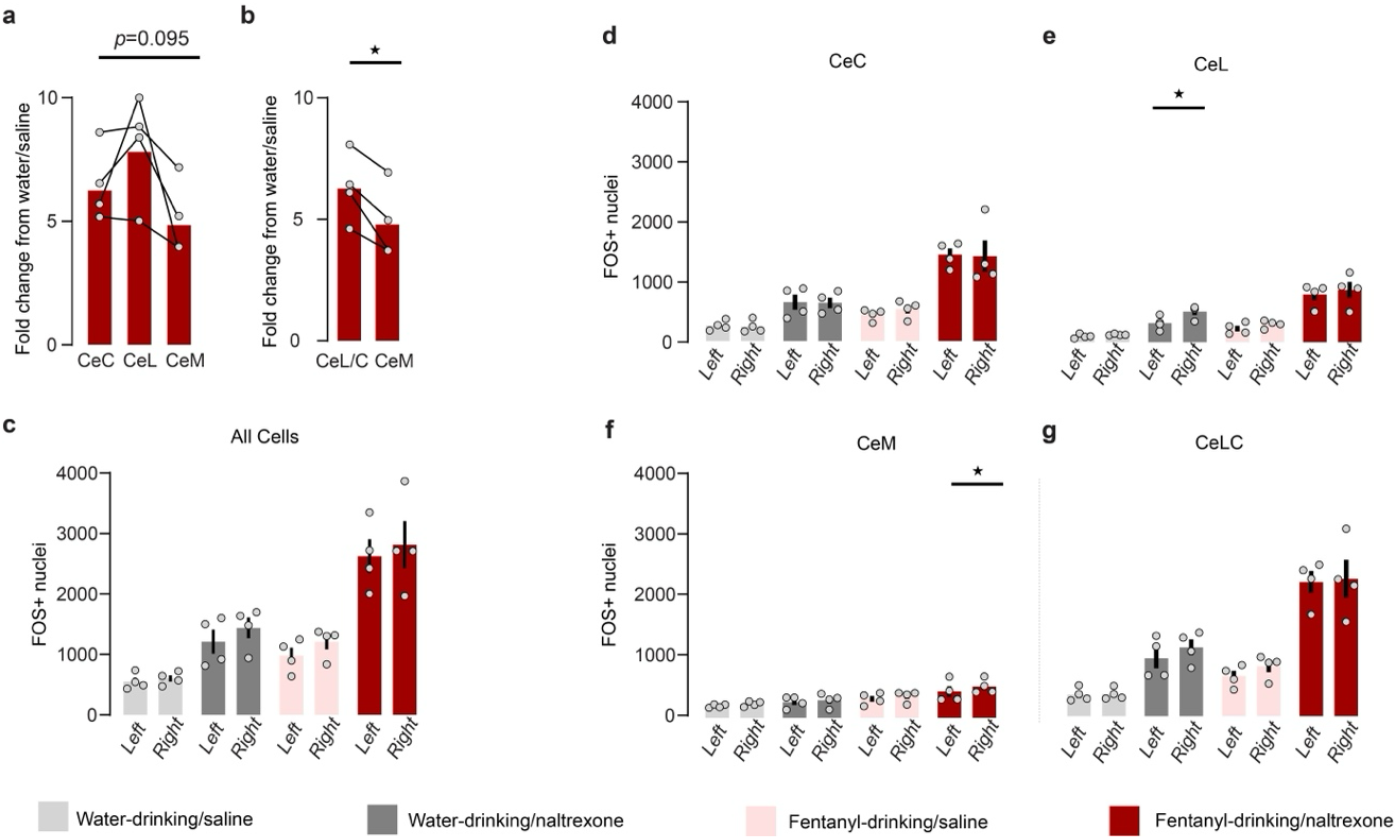
Further characterization of FOS^+^ nuclei in the CeA during fentanyl withdrawal. **(a)** Normalized fold-change in FOS^+^ nuclei in the fentanyl-drinking/naltrexone group, compared to water-drinking/saline condition. (One-way repeated measures ANOVA). **(b)** When considering the CeLC as a whole, we detected a greater increase in the number of FOS^+^ neurons in the CeLC compared to the increase in the CeM (Paired t-test, t=4.554, *p=0.0198). **(c)** We detected a main effect of hemisphere on the expression of FOS (top left; Three-way repeated measures ANOVA, main effect of side (F(1,12)=8.238, p=0141), drinking condition (F(1,12)=35.74, p<0.0001), and withdrawal condition (F(1,12) = 23.15, p=0.0004), but no comparisons were significant after Bonferroni’s correction. Bars represent mean value +/-SEM; grey dots represent individual subject values. **(e)** No main effect of hemisphere on the numbers of FOS^+^ nuclei in the CeC (Three-way repeated measures ANOVA, main effects of drinking condition (F(1,12)=25.32, p=0.0003), withdrawal condition (F(1,24)=43.17, p<0.0001), and drinking x withdrawal interaction (F(1,12)=7.405, p=0.0186). **(e)** We found more FOS^+^ nuclei in the right CeL compared to the left CeL in water-drinking mice after naltrexone administration (Three-way repeated measures ANOVA with Bonferroni’s correction, main effect of hemisphere (F(1, 12)=13.31, p=0.003), drinking condition (F(1,12) = 19.95, p=0.0008), and withdrawal condition (F(1,12)=46.86, p<0.0001); *p=0.0102). **(f)** We found more FOS^+^ nuclei in the right CeM compared to the left CeM in fentanyl-drinking mice after naltrexone administration (Three-way repeated measures ANOVA with Bonferroni’s correction, main effect of hemisphere (F(1, 12)=14.18, p=0.002) and drinking condition (F(1,12) = 19.864, p=0.0008). *p=0.0281. **(g)** No main effect of hemisphere on the numbers of FOS^+^ nuclei in the CeLC as a whole (Three-way repeated measures ANOVA, main effect of drinking condition (F(1,12)=30.33, p=0.0001), withdrawal (F(1,12)=58.35, p<0.0001), and drinking x withdrawal interaction (F(1,12)=8.084, p=0.0148).

**Supplementary Fig. 2.**
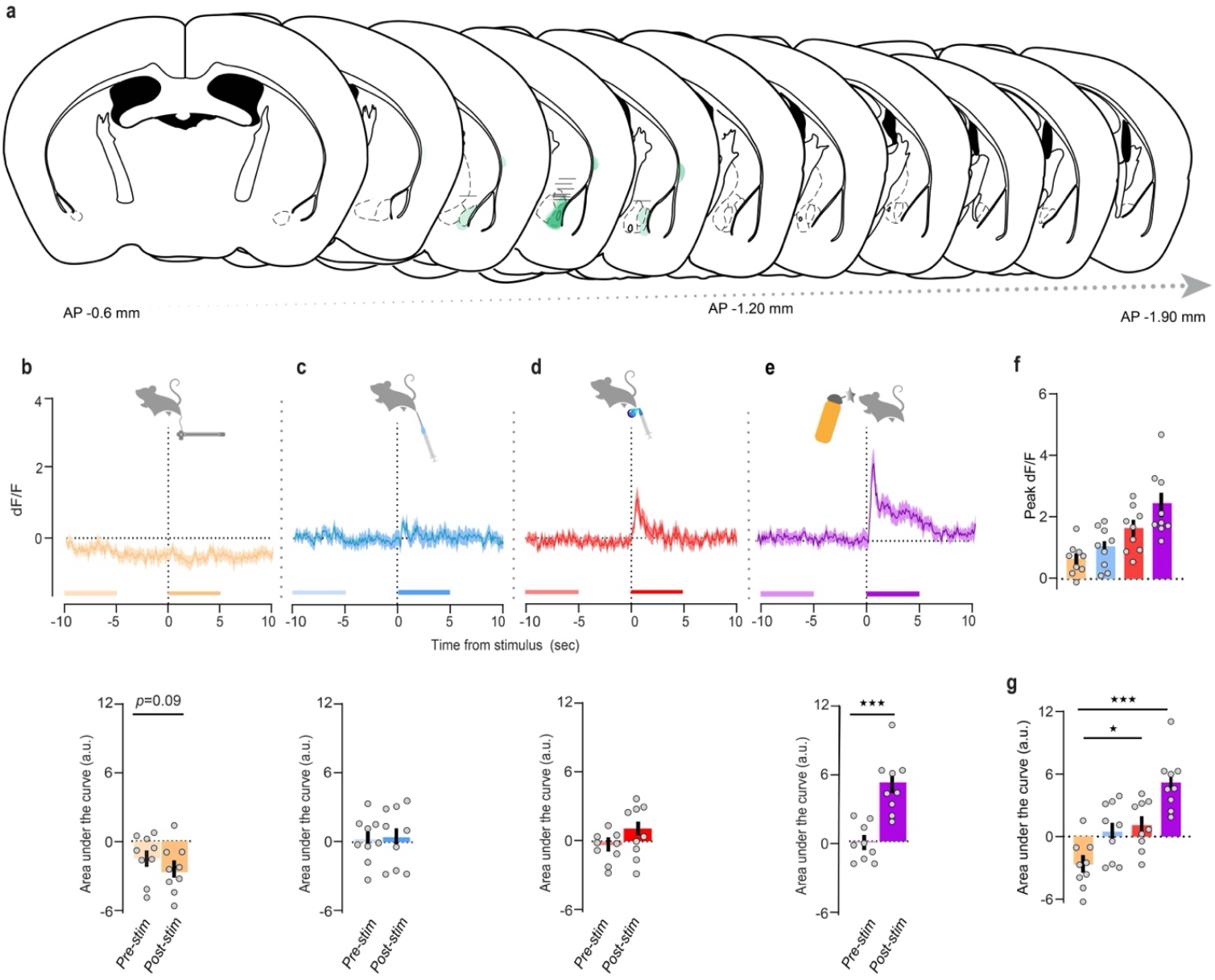
CeLC^PKC^^δ^ neurons in opioid-naïve mice respond to noxious aversive and non-noxious aversive stimuli. **(a)** Approximate location of fiber tip, and viral expression on the same slice, for each subject **(b)-(e)** Top: peri-stimulus time histogram of non-normalized dF/F (z-score) from −10sec to 10sec from the moment of stimulus application, following application of **(b)** a 0.16g von Frey filament (innocuous light touch); **(c)** noxious pin-prick with a 25G needle (noxious pinprick); **(d)** a 55°C hot water drop (noxious hot water); and **(e)** an aversive, but non-noxious, airpuff delivered to the side of the face contralateral to virus injection and fiber. Lines and area fill represent mean dF/F of 10 trials/subject, averaged across all subjects, +/-SEM. Bottom: area under the dF/F curve (A.U.C) prior to (−10 sec to −5 sec) vs. after (0 sec to 5 sec) each stimulus. **(b)** Compared to pre-stimulus baseline, innocuous light touch was associated with a decrease in the AUC, but this observation did not reach significance (Paired t-test, t=2.068). Bars represent mean dF/F +/-SEM; dots represent individual points. **(c)** Noxious pinprick and **(d)** noxious hot water did not significantly affect area under the dF/F curve. **(e)** Aversive airpuff was associated with a significant increase in area under the dF/F curve (Unpaired t-test, t=5.088, \*\**\*p*=0.0007). **(f)** Similar trends were observed between non-normalized dF/F and normalized z-scored dF/F in terms of peak responses to each stimuli, but this did not reach statistical significance (One-way repeated measures ANOVA). **(g)** Compared to innocuous light touch, aversive airpuff and hot water produced a significant increase in the AUC (One-way ANOVA with repeated measures and Bonferroni’s correction; significant effect of treatment (F(2.220, 17.76)=13.11, p=0.0002); *p=0.0135, ***p=0.0006; pinprick AUC was slightly raised, but it did not reach statistical significance (p=0.0549)

**Supplementary Fig. 3.**
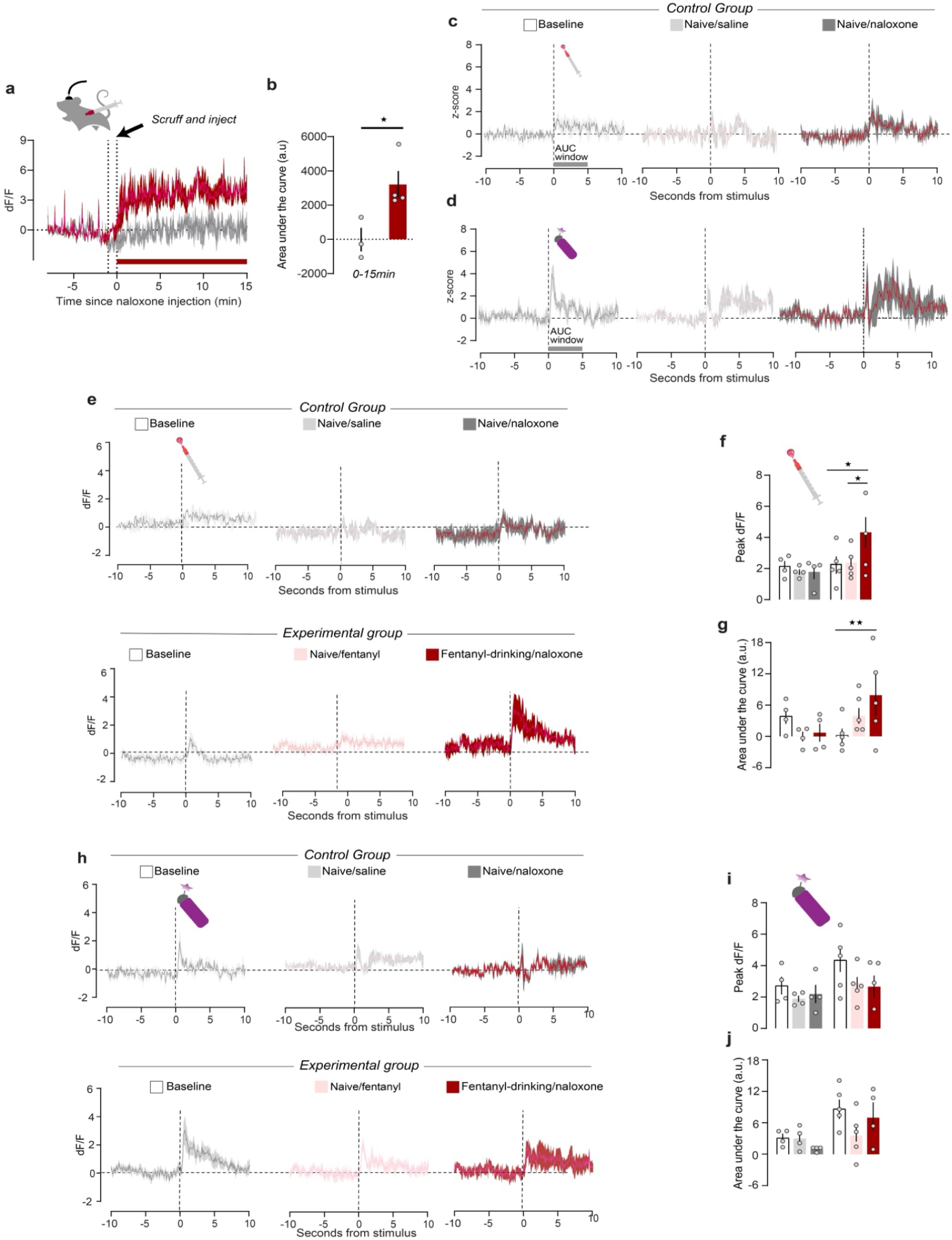
CeLC^PKCδ^ neurons are hyperactive and hypersensitive during fentanyl withdrawal. **(a)** Peri-injection time histogram of fluorescence before, during, and after an injection of 3 mg/kg naloxone in control, opioid-naïve animals (dark gray) and fentanyl-dependent animals (red). Red bar: AUC interval in **(b)**.Lines and area fills represent mean +/-SEM of the dF/F fluorescence for each timepoint; data were collected at 10Hz and downsampled to 3Hz during postprocessing. **(b)** The net AUC for the 15 min after naloxone injection is significantly higher for fentanyl-drinking vs. naïve mice (Unpaired t-test, t=2.951, *p=0.0319). **(c)-(d)** Peri-stimulus time histogram of the z-scored fluorescence from 10 sec prior to 10sec following application of a noxious hot water drop **(c)** or airpuff **(d)** for the water-drinking group on the baseline stimulus test day (left), after an injection of saline (middle), or after an injection of naloxone (right). Lines and area fill represent mean values of five trials averaged across subjects. **(e)** Peri-event time histogram of the dF/F fluorescence response to noxious hot water for the water drinking group (top) and fentanyl-drinking group (bottom) following each treatment condition. **(f)** Peak dF/F significantly increased following the application of a noxious hot water drop during withdrawal compared to the same animals at baseline or following 0.2 mg/kg fentanyl (Mixed-effects analysis with repeated measures and Bonferroni’s correction; significant Treatment x Timepoint interaction (F(2,13)=4.268; p=0.0376). *p=0.0138 (Baseline vs. fentanyl/naloxone) or p=0.019 (Acute fentanyl vs. fentanyl/naloxone). **(g)** The AUC of the dF/F curve for noxious hot water response curve was significantly higher during fentanyl withdrawal than at baseline (Mixed-effects analysis with repeated measures and Bonferroni’s correction, significant Treatment x Timepoint interaction (F(2,13)=6.572, p=0.0107; **p=0.0065). **(h)** Peri-event time histogram of the dF/F fluorescence response to airpuff for the water drinking group (top) and fentanyl-drinking group (bottom) following each treatment condition. **(i)** There was no effect of treatment group or experimental timepoint on the peak dF/F fluorescence response to an aversive airpuff (Mixed-effects analysis with repeated measures). **(j)** There was no effect of treatment group or experimental timepoint on the AUC of the dF/F curve an aversive airpuff (Mixed-effects analysis with repeated measures)

**Supplementary Fig. 4.**
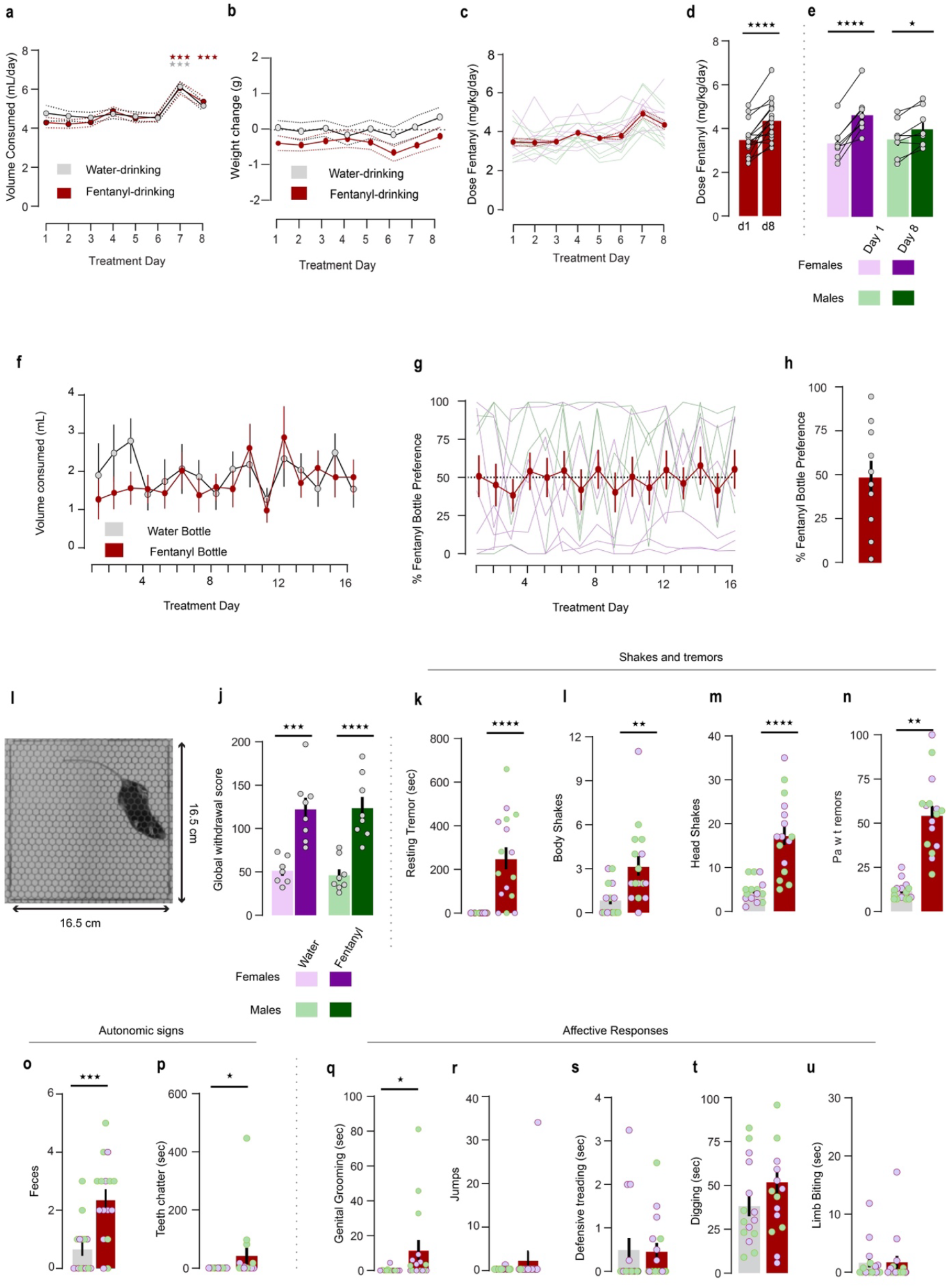
Further characterization of fentanyl dependence phenotype. **(a)-(e)** Patterns of fentanyl drinking and intake. **(a)** Both untreated water-drinking mice (i.e., water-drinking mice; grey dots/line, n=7 female and 8 male mice) and fentanyl treated water-drinking mice (i.e., fentanyl-drinking mice; red dots/line, n=8 female and 8 male mice) consumed significantly more liquid on Treatment Day 7, and fentanyl-drinking mice consumed significantly more liquid on Treatment Day 8, compared to consumption on Treatment Day 1 (Two-Way Repeated Measures ANOVA, significant effect of Treatment Day; F(7, 203)=22.74, p<0.0001). There was also a significant effect of subject (F(29, 203)=11.50, p<0.0001) but no significant effect of drinking condition or Treatment Day x drinking condition interaction. Each point represents mean values, dotted lines are +/-SEM. ***p<0.001 compared to Treatment Day 1 (Bonferroni post-hoc test). **(b)** Fentanyl-drinking mice did not show significant weight changes from water-drinking mice on any treatment day (Mixed-effects analysis, main effect of Treatment Day (F(7, 195)=3.097) **(c)-(e)** *Prkcd*-Cre mice do not develop a significant preference for the fentanyl-treated water bottle in a two-bottle choice paradigm. n=6 females and 4 males **(c)** Fentanyl-drinking mice consumed 2.33-6.67 mg/kg/day fentanyl throughout the treatment period, trending with total fluid consumption. Points represent mean +/-SEM. **(d)** Fentanyl dose consumed on Treatment Day 8 was significantly higher than the dose consumed on Treatment Day 1 (Paired t-test, t = 5.917, ****p<0.0001). **(e)** A two-way ANOVA comparing intake on Treatment Day 1 vs. Treatment Day 8 for male and female mice found a main effect of treatment day (F(1,14)=68.27, p<0.0001), subject (F(14,14)=15.36, p<0.0001), and a sex x treatment day interaction (F(1,14)=15.24, p=0.0014), but no main effect of sex on the dose consumed. Female (purple) and male (green) mice consumed a higher dose on treatment day 8 (dark bar) than on treatment day 1 (light bar; ****p<0.0001, *p=0.0162). Bars represent mean dose consumed +/-SEM; grey dots represent individual subjects’ values. **(f)-(h)** Fentanyl drinking model does not support two-bottle choice. **(f)** Mice consumed similar volumes from the water and fentanyl bottle on every treatment day (Mixed-effects analysis). Points represent the mean +/-SEM volume consumed from each bottle on each day. **(g)** The preference for the fentanyl bottle vs. the water bottle did not increase over the course of the two-bottle choice paradigm (Mixed effects analysis) Points represent the mean +/-SEM dose consumed on each treatment day; lines represent individual mice’s trajectories (purple: female mice; green: male mice). **(h)** The mean preference scores for individual animals ranged from 2.32-94.88% preference for the fentanyl bottle over the water bottle. Bar represents mean % fentanyl bottle preference +/-SEM. **(i)** View of mouse during withdrawal scoring. Cameras mounted directly underneath the observation chamber were placed inside a Plexiglas box, allowed for a clear view of the animal’s jaw and paws. Videos were recorded under red light and desaturated to facilitate scoring. (**j)** Global withdrawal scores were similar between water-drinking male and female mice (light bars) and between fentanyl-drinking male and female mice (dark bars; Two-way ANOVA, main effect of treatment (F(1,27)=48.10, p<0.0001; ***p=0.0002, ****p<0.0001). **(k)-(u)** Individual measures that contribute to global withdrawal score. **(k)** Fentanyl-drinking mice spent significantly more time in a resting tremor vs. water-drinking mice (Mann Whitney test, U=22.50, ****p<0.0001). Purple points: female mice; green points: male mice **(l)** Fentanyl-drinking mice exhibited significantly more body shakes vs. water-drinking mice (Mann Whitney test, U=43.50, **p<0.0014). **(m)** Fentanyl-drinking mice exhibited significantly more head shakes vs. water-drinking mice (Unpaired t-test, t=5.386, ****p<0.0001). **(n)** Fentanyl-drinking mice exhibited significantly more paw tremors vs. water-drinking mice (Unpaired t-test, t=3.096, **p=0.0043). **(o)** Fentanyl withdrawal was associated with increased feces expelled compared to water-drinking mice (Unpaired t-test, t=3.994, ***p=0.0004). **(p)** Fentanyl-drinking mice exhibited significantly more teeth chattering vs. water-drinking mice (Mann Whitney test, U=75, *p=0.0177). **(q)** Fentanyl withdrawal increased time spent engaging in genital grooming (Mann Whitney test, U=66, *p=0.0143) However, fentanyl withdrawal was not associated with an increase in **(r)** jumps (Mann Whitney Test) or **(s)** time spent defensive treading (Mann Whitney test), **(t)** Digging (Unpaired t-test), or **(u)** Limb biting (Mann Whitney test).

**Supplementary Fig. 5.**
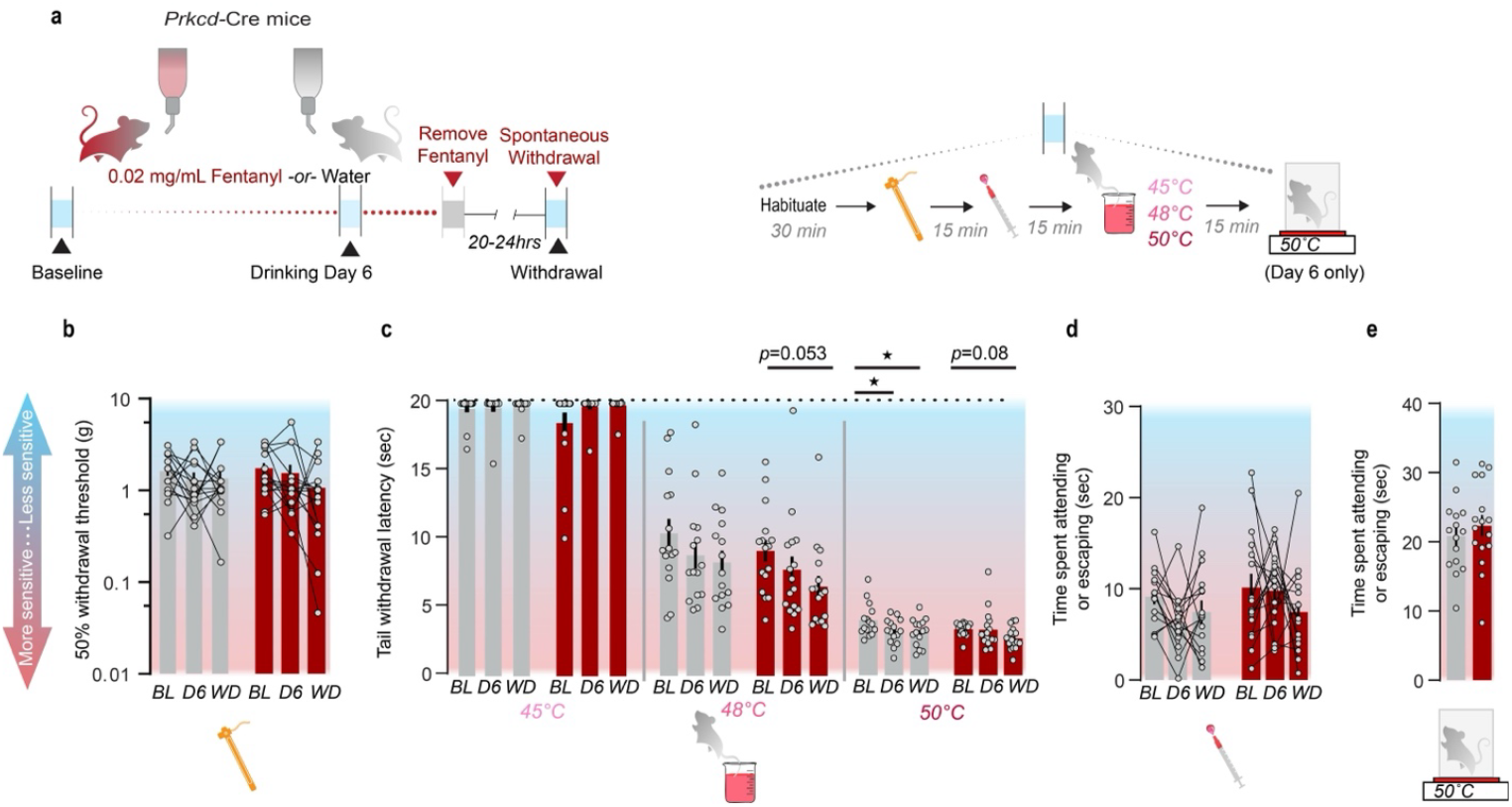
Fentanyl exposure in home-cage water supply does not induce hyperalgesia during the course of treatment or during spontaneous withdrawal. **(a)** Left: Experimental timeline. Test days are indicated with grey bars and black arrows. Right: outline of procedure on testing days. Because mice rapidly learn to engage in escape behaviors when repeatedly placed on an inescapable hotplate, we only tested mice on the hotplate on Treatment Day 6. **(b)-(c)** No effect of fentanyl treatment on reflexive responses to tactile or thermal stimuli. **(b)** Six days of fentanyl treatment and spontaneous withdrawal had no effect on 50% withdrawal threshold in the von Frey Up-Down assay (Two-way Repeated Measures ANOVA). Grey bars: water-drinking mice; red bars: fentanyl-drinking mice. Bars represent mean values +/-SEM; points represent individual subjects’ data. Blue shading indicates less sensitivity to stimuli (i.e., higher nociceptive thresholds); red shading indicates more sensitivity (i.e., lower nociceptive thresholds). (**c)** Fentanyl had no effect on mice’s latency to withdraw their tails from 45°C (left), 48°C (middle), or 50°C water (right). 45°C: Two-way repeated measures ANOVA, significant effect of subject only (F(29, 58)=2.190, p=0.0056). 48°C: Two-way repeated measures ANOVA with Bonferroni’s correction, main effect of timepoint (F(2,58)=4.185, p=0.02) and subject (F(29,58)=1.674, p=0.0476). 50°C: Two-way repeated measures ANOVA with Bonferroni’s correction, main effect of timepoint (F(2, 58)=5.520, p=0.0064). We detected no main effect of treatment group or an interaction on tail withdrawal latency at any temperature. *p<0.05. **(d)-(e)** No effect of fentanyl treatment on affective responses to noxious thermal stimuli. **(d)** No effect of fentanyl or experimental timepoint on the time spent engaging in attending or escape responses to a noxious hot water droplet applied to the left hindpaw. (Two-way repeated measures ANOVA.) **(e)** No effect of fentanyl treatment on the time spent engaging in attending or escape behaviors on an inescapable 50°C hotplate. (Two-way repeated measures ANOVA).

**Supplementary Fig. 6.**
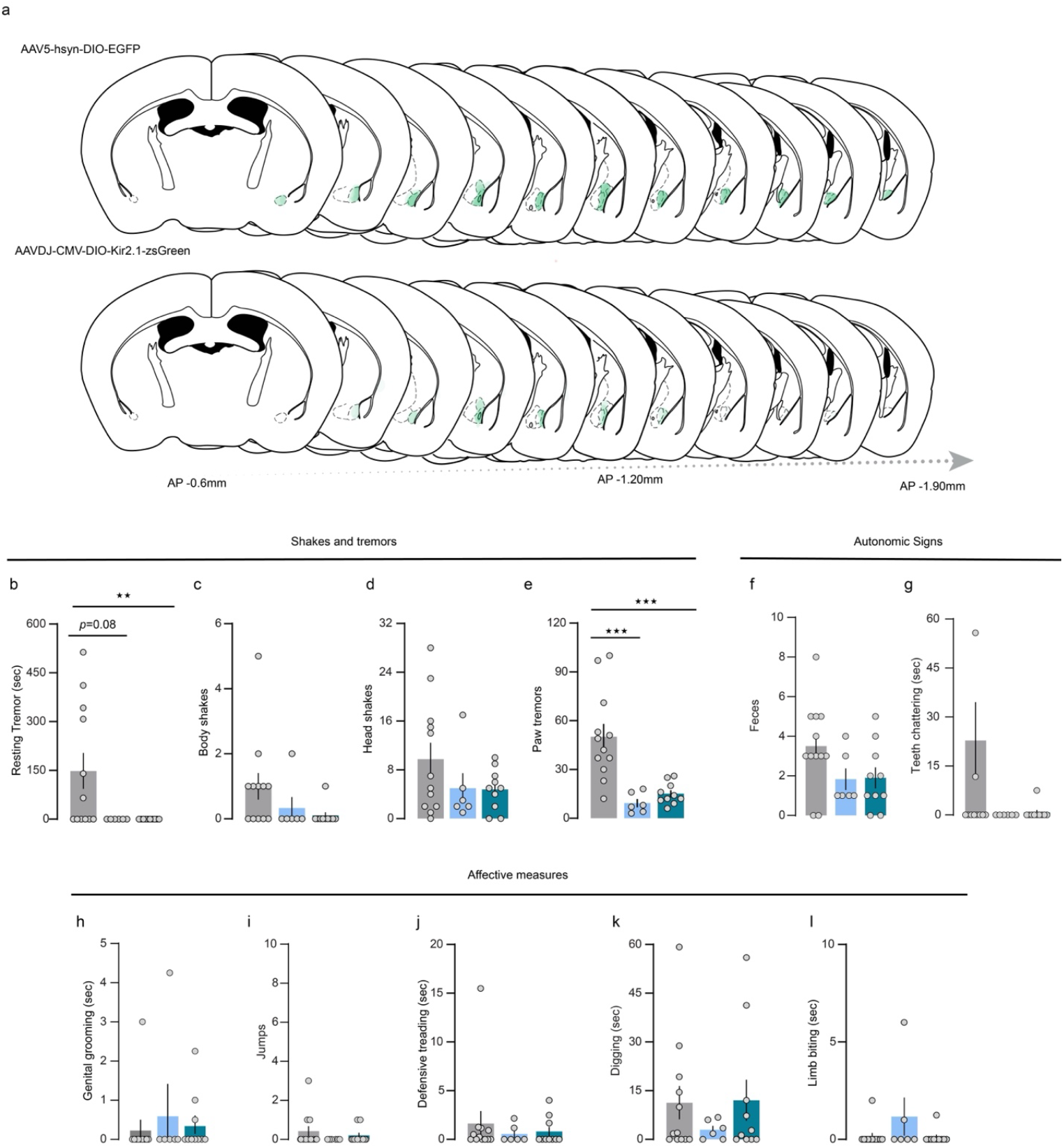
Further characterization of Kir2.1 overexpression-suppressed withdrawal phenotype. **(a)** Viral spread for individual subjects. Top: AAV5-hsyn-DIO-EGFP viral spread (n=6 females and 6 males. Bottom: AAVDJ-CMV-DIO-Kir2.1-zsGreen viral spread. n=9 females and 7 males. **(b)** Kir2.1 overexpression significantly decreased the time with a resting tremor in fentanyl-dependent mice (One-way ANOVA, F(2,25) =4.666, p=0.019, *p=0.0347). Kir2.1 overexpression did not result in fewer **(c)** body shakes or **(d)** head shakes in fentanyl-drinking mice (One-way ANOVA). **(e)** Kir2.1 overexpression significantly decreased the number of paw shakes in fentanyl-dependent mice (F(2,25)=13.79, p<0.0001, ***p=0.0006). Kir2.1 overexpression did not significantly alter **(f)** the number of feces expelled, **(g)** time with teeth chattering, **(h)** time engaged in genital grooming, **(i)** jumps, **(j)** time spent defensive treading, **(k)** time spent digging, or **(l)** limb biting (One-way ANOVAs).

**Supplementary Fig. 7.**
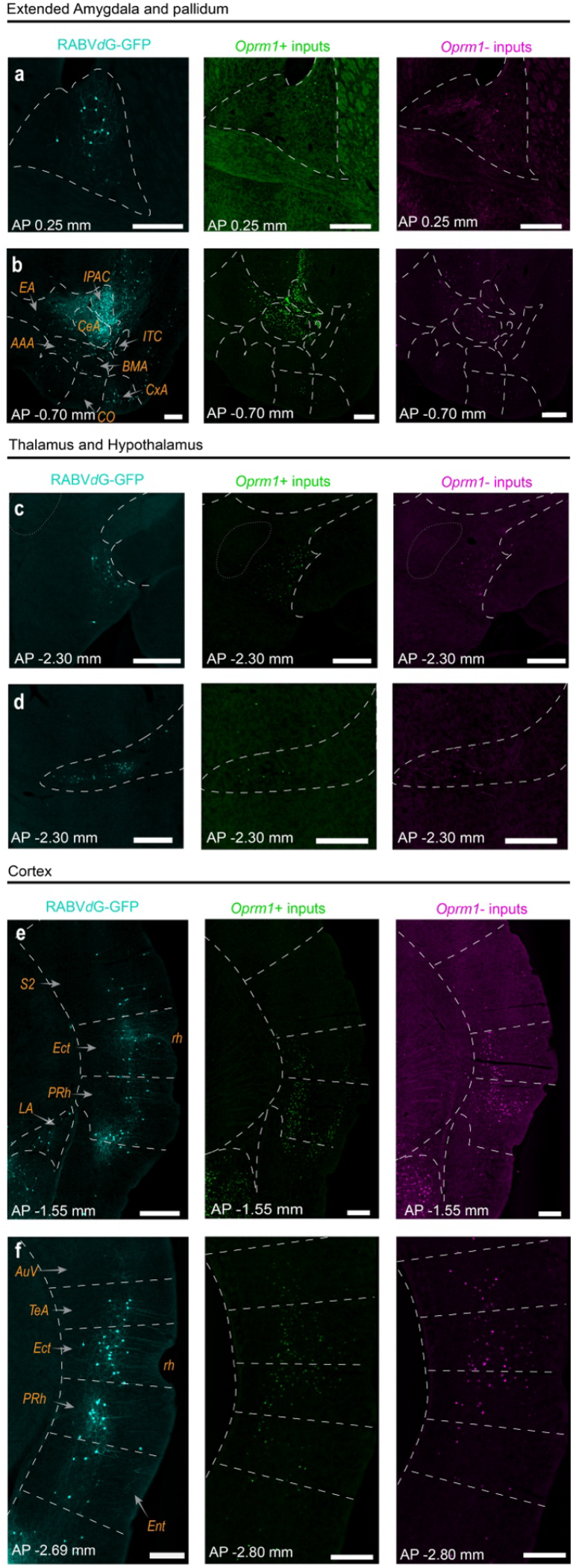
Opioid-sensitive neurons in brain regions containing putative monosynaptic inputs to CeLC^PKCδ^ neurons. **(a)**-**(b)** RABV*d*G (cyan), EGFP (i.e., *Oprm1*^+^ cells; green) or mCherry (i.e., *Oprm1*-cells; magenta) labeling in the anterior extended amygdala and nearby structures. **(a)** Labeling in the bed nucleus of the stria terminalis (BNST), particularly the oval nucleus. Scale bars: 250 um. **(b)** Labeling in the anterior amygdala, extended amygdala, olfactory, and nearby regions. Scale bars: 250 um. Abbreviations: AAA: anterior amygdaloid area; BMA: basomedial amygdala; CeA: CeA; CO: Cortical amygdala; CxA: Cortex-amygdala transition zone; EA: extension of the amygdala; END: endopiriform nucleus; ITC: intercalated nuclei of the amygdala. **(c)-(d):** Labeling in the thalamus and hypothalamus. **(c)**: labeling in the lateral hypothalamus (LH), particularly in the parasubthalamic nucleus. Scale bars: 250µm. **(d)** Labeling in the ventral posterior nucleus of the thalamus (VENT). Scale bars: 250 um. **(e)-(f):** Labeling in the cortex. **(e)** Labeling in the medial portions of the temporal cortex. Abbreviations: ECT: ectorhinal cortex; PRh: perirhinal cortex; rh: rhinal fissure; S2: secondary somatosensory area, LA: Lateral amygdala. Scale bars: 250um. **(f)** Labeling in the posterior temporal cortex. Abbreviations: Au1: primary auditory cortex; ENT: entorhinal cortex; TeA: Temporal association cortex. Scale bars: 250um.

**Supplementary Fig. 8.**
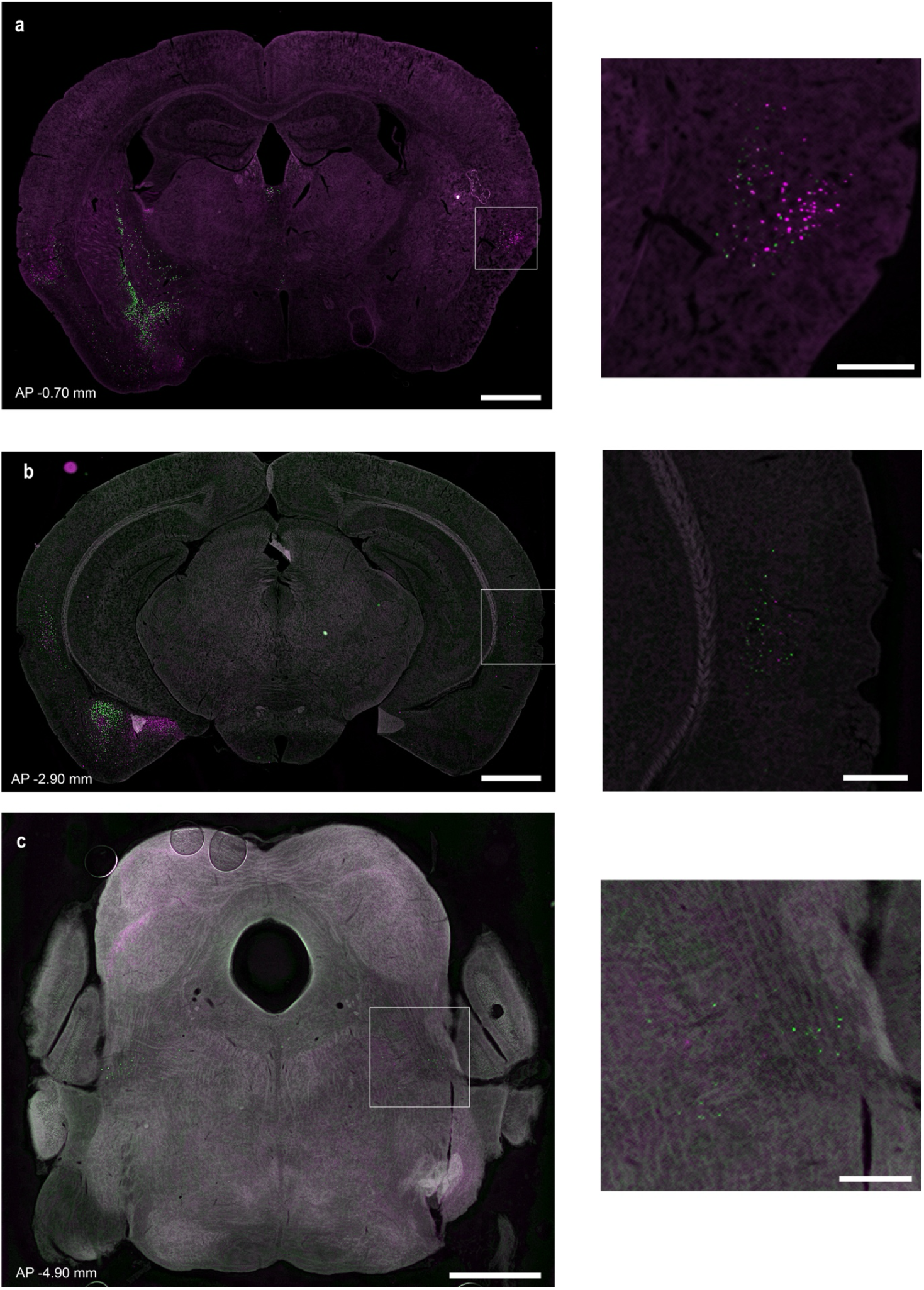
Sparse labeling of contralateral *Oprm1*^+^ and *Oprm1*^-^ inputs to the CeA with a retrograde, *Oprm1*^+^/*Oprm1*^-^ neuron-labeling virus. **(a)** Labeled nuclei are visible bilaterally in the contralateral insular cortex, but not in the striatal and extended amygdala areas with extensive ipsi-lateral inputs. Scale bars for all panels: Large image,1 mm; inset, 250 um. **(b)** Labeled nuclei are visible bilaterally in the cortical areas adjacent to the rhinal fissure, but not in the posterior BLA or the amygdalohippocampal area that show dense ipsilateral inputs. **(c)** Labeled nuclei are visible bilaterally in the parabrachial nucleus.

**Supplementary Fig. 9.**
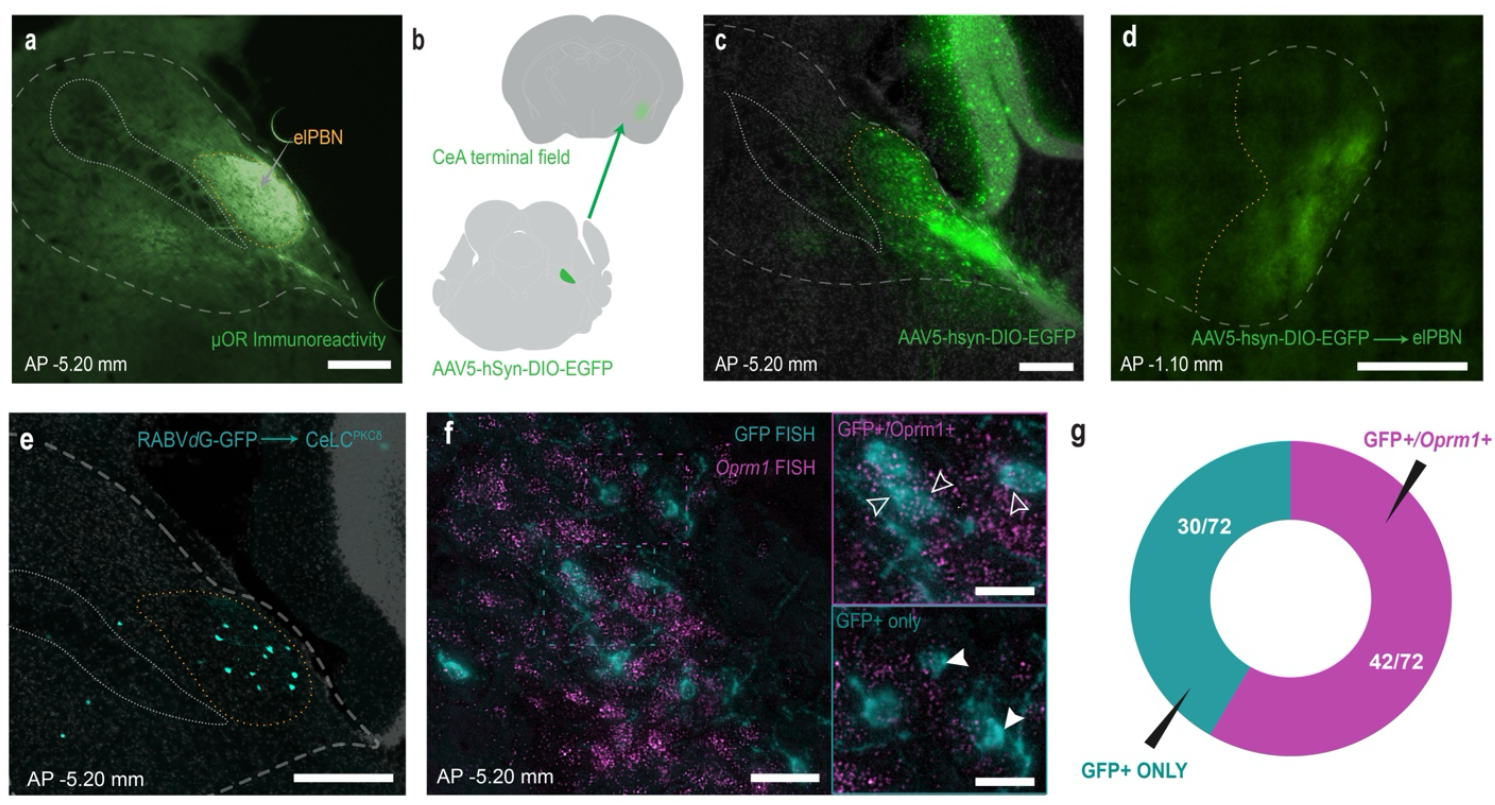
Opioid-sensitive neurons in the parabrachial nucleus project to CeLC^PKCδ^ neurons. **(a)** MOR immunoreactivity in the parabrachial nucleus, especially its external lateral nucleus (elPBN; orange). Scale bars: 250 um **(b)** Experimental design. We injected the elPBN of *Oprm1*-Cre mice with AAV5-hsyn-DIO-EGFP, which allowed us to visualize *Oprm1+* cell bodies in the parabrachial nucleus (PBN^MOR^) and their terminals in the CeA. **(c)** Cre-dependent EGFP expression in the parabrachial nucleus of an *Oprm1*-Cre mouse. Scale bars: 250 um **(d)** PBN^MOR^ terminals are restricted to the capsular CeC in the anterior part of the CeA. **(e)** Expression of RABVdG-GFP (Cyan) in the elPBN. We injected Cre-dependent helper AAVs, followed by RABV*d*G-GFP, into the CeA of *Prkcd*-Cre mice. Scale bars: 250 um. **(f)** *Oprm1* mRNA (magenta) colocalizes with RABV*d*G-GFP (cyan) expression in the elPBN of *Prkcd*-Cre mice, suggesting that PBN^MOR^ neurons send monosynaptic inputs to CeLCPKd neurons. Scale bar: 250 um; insets: 100 um. Filled arrowheads: GFP+/*Oprm1*-neurons; open arrowheads: GFP+/*Oprm1*^+^ neurons **(g)** quantification of **(f)**. n=5 mice, 5 slices/mouse, 72 identified GFP+ cells after RNAScope.

## Notes

### Competing Interest Statement

The authors have declared no competing interest.

